# Transcriptomic shift in ethanol and amino acid metabolic genes regulated by Med15 during alcoholic fermentation

**DOI:** 10.64898/2026.01.08.698492

**Authors:** David G. Cooper, Emma Grunkemeyer, Jan S. Fassler

## Abstract

Organisms that thrive in extreme environments provide natural experiments in evolution, revealing the genetic regulators that orchestrate complex phenotypic change. Wine yeast are specialized strains that are adapted to survive in the wine making environment while producing high concentrations of ethanol. In addition to large genomic changes that differentiate wine yeast from yeast used in other industries, single nucleotide and polyglutamine tract polymorphisms in the transcriptional regulator Med15 are associated with the fermentation efficiency and stress response phenotypes of wine yeast. In this study we investigated the transcriptional differences during wine fermentation in transgenic lab strain yeast having integrated wine yeast *MED15* alleles. Compared to the unmodified lab strain (LAB or *MED15*_LAB_), the same strain in which the *MED15* locus was replaced with a *MED15* allele from yeast isolated from palm wine, the fermented sap of palm (oil, date, coconut) trees, (WY23, or *MED15*_WY23_) exhibited enhanced expression of glycolytic, fermentation, and amino acid biosynthesis genes. Our experimental data confirms the importance of arginine biosynthetic genes during the fermentation process and suggests that the improvement in fermentation efficiency in strains with *MED15* alleles from some wine yeast strains may be related to the role of Med15 in expression of the genes of the arginine biosynthetic pathway. The global benefit conferred by polymorphisms in a single transcriptional regulator, makes Med15 a prime target for engineering of strains devoted to various types of alcohol production.

## Introduction

The production of wine requires yeast strains that can efficiently convert sugars to ethanol while also surviving the stresses of the grape juice/must environment including high osmolarity, low pH, low nitrogen availability, and ultimately high ethanol concentrations. The general transcriptional regulator Med15 influences both fermentation efficiency and stress response during grape juice/must fermentation (1) and thus the *MED15* gene is a good candidate for domestication niche-specific adaptative mutations leading to improved fermentation observed in wine yeast (1, 2). Variations in alleles of the general transcriptional regulator *MED15*, including the length of several polyglutamine (poly-Q) tracts embedded in Med15 sequence, are associated with yeast domestication niches (beer and wine) (2).

While various yeast species are naturally present on grapes, *S. cerevisiae* dominates the fermentation environment (3). Historically, and as a modern trend, uninoculated “natural fermentations” have been used in the production of wine, though many wineries will specifically inoculate grape must with previously isolated and commercially available wine yeast strains (4).

Fermentation reactions are characterized by an initial lag phase for yeast not pre-adapted to juice followed by a growth phase after which cells reach saturation (10^8^ cells/mL). Available nitrogen and ∼50% of the sugars are quickly consumed during the growth phase (5). The remaining sugar is fermented after culture saturation. During this time, the sugar is used almost exclusively to generate energy and survival compounds rather than biomass (5). Ethanol is the main byproduct of yeast fermenting glucose for energy. Other metabolic products are made throughout the fermentation process including various acids (acetic, succinic, and lactic) and volatile compounds (esters, fatty acids, aldehydes, and higher alcohols) (6–8). Industrial fermentation reactions can be reliably replicated in the lab using small scale fermentation reactions (1, 8).

In addition to environmental stresses such as high sugar content osmotic stress, nitrogen limitation, and low pH present at the start of fermentation, accumulation of both ethanol and acetic acid are stressors specific to late fermentation cultures (3). Correspondingly, wine yeast exhibit high tolerance to many compounds and stressors including acetic acid, copper, sulfite, high glucose, sorbitol, and ethanol (9). Yeast respond to the initial and accumulating stresses of grape juice and fermentation through the Environmental Stress Response (ESR) (10, 11). The ESR allows yeast to deal with fermentation stress and moderate levels of ethanol but eventually the levels of alcohol become toxic. Ethanol toxicity is thought to occur by disrupting membrane structural integrity, while the addition of lipids or oxygenation, that is required to synthesize fatty acids, can mitigate the toxic effects (6, 7).

The *S. cerevisiae* yeast strains used in food and beverage production (*e.g*., wine, beer, and bread) are phenotypically distinct from wild yeast strains and represent lineages from a small set of common ancestors (9, 12). Prior to the isolation of specific wine yeast strains, which are now removed from evolutionary processes by being maintained as stocks and sold to winemakers to add to grape must (13), wine yeast diverged from wild yeast populations in a limited number of domestication events (4, 9, 14).

Across different alcoholic beverage industrial niches, *S. cerevisiae* yeast are critical for their ability to efficiently ferment various carbon sources to produce ethanol and carbon dioxide. Yeast will preferentially undergo alcoholic fermentation and accumulate ethanol even if oxygen is readily available for respiration (3), a phenomenon known as the Crabtree effect (15, 16). After glycolysis, pyruvate is metabolized into acetaldehyde by Pyruvate DeCarboxylases (PDCs) and acetaldehyde is converted into ethanol by Alcohol DeHydrogenases (ADHs). This step serves to restore NAD^+^ by oxidation of NADH that is produced during glycolysis. While ancestral ADH genes were optimized to produce ethanol, the whole genome duplication within the clade that includes *S. cerevisiae* resulted in ADH genes encoding enzymes with the ability to more efficiently consume ethanol instead of exclusively producing ethanol (3). Specifically, Adh2 preferentially consumes ethanol with a Km for ethanol (600-800 μmol/L) that is significantly lower than that for Adh1 (17,000-20,000 μmol/L) (17, 18). Thus Adh2 works much more efficiently than Adh1 at oxidizing ethanol while the remaining four Adh proteins (mitochondrial Adh3 and Adh4, cytoplasmic Adh1 and Adh5) preferentially produce ethanol (17). Acetaldehyde produced by Adh2 is metabolized into acetate by ALdehyde Dehydrogenases (ALDs) and acetate is metabolized into Acetyl CoA by Acetyl CoA Synthetases (ACSs) to be used in the citric acid cycle or other metabolic pathways.

The loss of cis-regulatory signals for respiration genes in yeast following the whole genome duplication may have contributed to the Crabtree effect (19). Specifically, yeast diverging after the whole genome duplication, including *S. cerevisiae*, lost regulatory signals that would promote the expression of mitochondrial genes causing these yeast to display faster anaerobic growth compared to aerobic growth (19). The ability for *S. cerevisiae* to preferentially ferment sugars and to tolerate higher levels of ethanol was likely selected for as these traits allow *S. cerevisiae* to outcompete other yeast species in natural fermentation environments (3).

As a general transcriptional regulator, Med15 has global effects on stress responses and metabolism. Changes in the sequence of Med15 that influence activity could thus produce broad phenotypic changes which provide the opportunity for yeast to adapt to novel environments. Variation in *MED15* sequence across strains predominantly consists of small changes in the length of simple sequence repeats that underlie glutamine and glutamine-alanine tracts. Recently the phenotypic stress response consequences of Med15 polyglutamine variation have been investigated (1, 20, 21). Specifically, Med15 was found to be required for resistance to the coal-cleaning chemical, 4-methylcyclohexane methanol (MCHM), and changes in poly-Q tract lengths were found in yeast resistant to MCHM relative to the endogenous *MED15* allele in the lab strain (20). Our previous work revealed that some of the fermentation efficiency characteristic of wine yeast strains is conferred by SNPs and polyglutamine tract length polymorphisms in Med15 (1). We have found that the disorder promoted by glutamine residues is important for Med15 function and that variation in polyglutamine tract length in Med15 can modulate stress response and metabolic activity (21). The significance of polyglutamine tract length variation has also been investigated in the yeast Cyc8 transcriptional co-repressor protein (22) as well as in the ANGUSTIFOLIA and Clock proteins in other organisms where even small changes in tract length (2-4 additional glutamine residues) are associated with noticeable phenotypic changes (23–25).

Here we conduct, to our knowledge, one of the first analyses of the *MED15* transcriptome during fermentation to identify *MED15* regulated genes that are unique to the fermentation environment compared to standard lab conditions. In addition, we test the hypothesis that the *MED15* alleles from wine yeast harboring variable length polyglutamine tracts and other dispersed SNPS (1, 2) may have contributed to specialized fermentation transcriptomes by comparing the fermentation transcriptome of LAB strains into which we’ve introduced wine yeast MED15 alleles (labeled *MED15*_WY##_ or WY##) to the unmodified LAB strain (*MED15*_LAB_ or LAB) background. Briefly our results show that *MED15* regulates glycolytic and fermentation pathways, with a particularly strong and distinctive regulatory effect during fermentation and that some transplanted *MED15_WY_* alleles cause subtle but biologically significant changes in metabolism. We report the transcriptomic basis for the improvement in fermentation rates seen in specific alcoholic beverage yeasts previously characterized (1) and although we expected changes to glycolysis and ethanol metabolism gene expression due to the *MED15*_WY_ alleles, we found a host of other differences in metabolic gene expression relative to unmodified laboratory strains (*MED15_LAB_*) suggesting that there are many different transcriptomes that can enhance fermentation rates.

## Methods

### Strains and Plasmids

Strains used in this study are derivatives of S288C, a widely used non-flocculent laboratory strain, originally designed for biochemical studies (26). The *med15* deletion strain is from the deletion collection (27). Genotypes of additional strains used in this study are listed in Table 1. Wine yeast (WY) strains were obtained from the ARS culture collection (https://nrrl.ncaur.usda.gov/) (WY23, palm wine yeast, NRRL-Y17772) or were from the Goddard laboratory collection (28) (WY7, Lalvin ICV D-254; WY15, DSM Fermichamp). Plasmids used in this study are listed in Table 2.

**Table 1.**
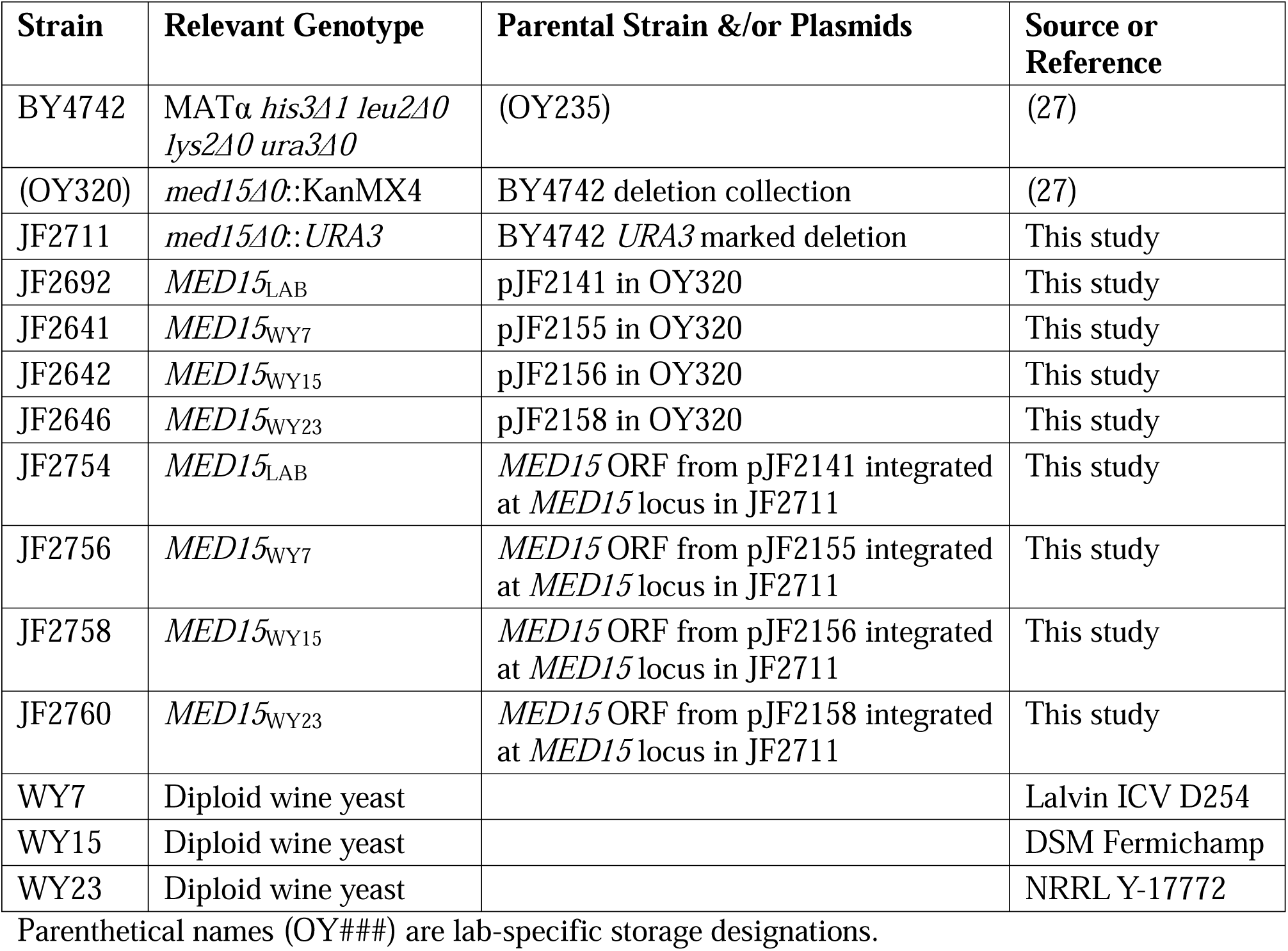
Strains.

**Table 2.**
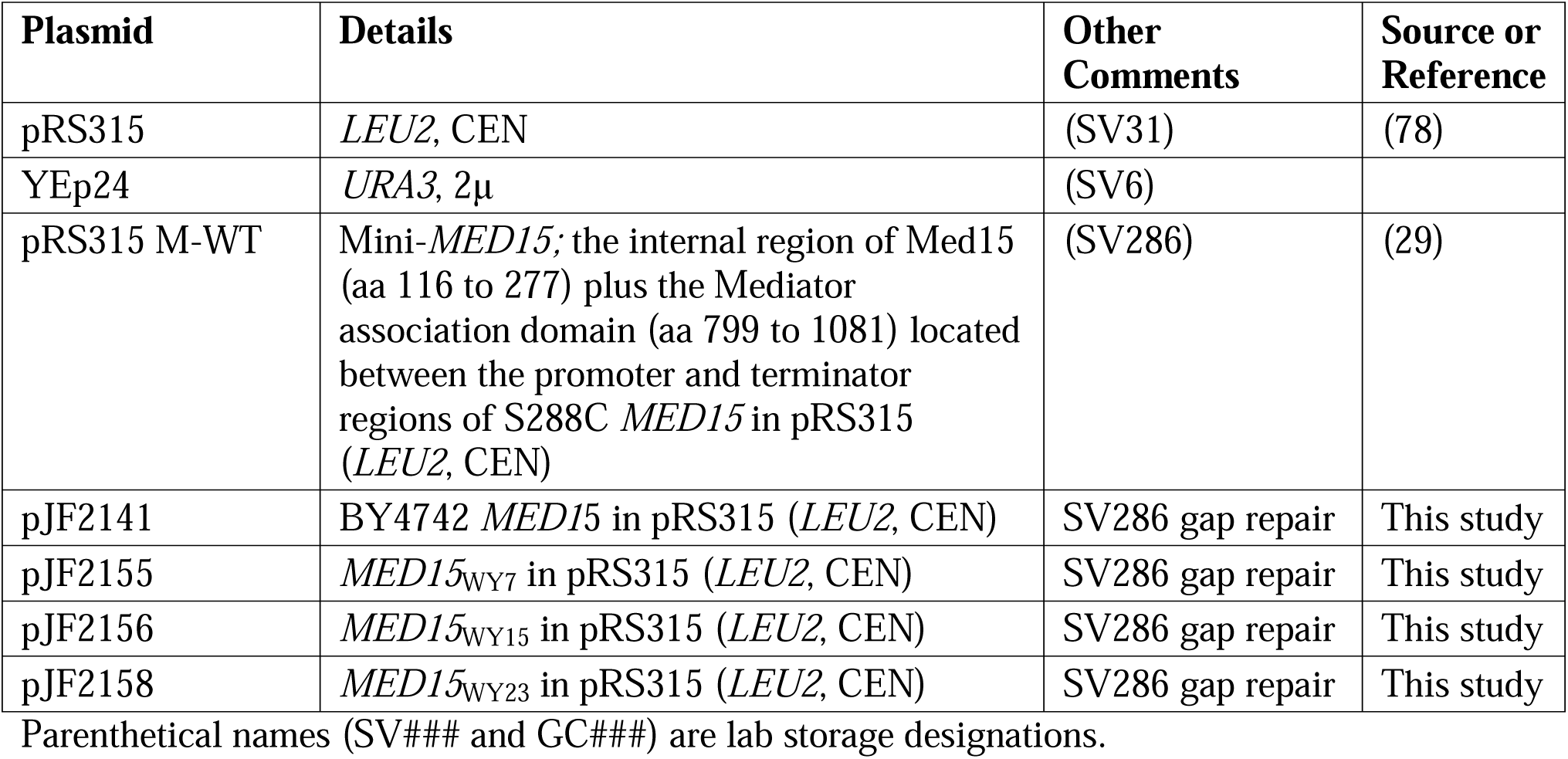
Plasmids.

### Wine Yeast *MED15* Integration

A *URA3* marked *med15Δ* lab strain (JF2711) was created to use as a starting point for integrating wine yeast *MED15* alleles. A *URA3* expression cassette was amplified from YEp24 using *URA3* GR (gap repair) F1 and R1 primers. Primers were complementary to sequences outside of the *MED15* ORF, allowing for homology to be used in the gap repair of *Bam*HI and *Spe*I digested Mini-*MED15* plasmid (pRS315 M-WT) (29). The *URA3* expression cassette with *MED15* flanking homology was subsequently amplified from the newly constructed plasmid using *MED15* F-245 and *MED15* R+3498 primers and integrated into the genome of BY4742 at the *MED15* locus to generate JF2711. Wine yeast *MED15* alleles were integrated into the *MED15* locus of JF2711 by homologous recombination. *MED15* coding sequences with 250 bp of upstream and downstream sequence were amplified from expression plasmids (pJF2141, LAB; pJF2155, WY7; pJF2156, WY15; pJF2158, WY23) using *MED15* F-245 and *MED15* R+3498 primers. *MED15* PCR fragments were then transformed into JF2711 and cells plated on YPD. After a day of growth, plates containing 800 mg/L 5-fluoroorotic acid (5-FOA) were used to counter-select transformants that had become Ura- due to integration of the new allele and loss of the *URA3* marked *MED15* locus. Integrants were stored as JF2754-JF2760.

### Colony PCR

Yeast colonies were used for rapid PCR screening using Taq polymerase, colony PCR buffer (final concentrations: 12.5 mM Tris-Cl (pH 8.5), 56 mM KCl, 1.5 mM MgCl2, 0.2 mM dNTPs), and primers at a final concentration of 0.2 μM. Reaction mixes were aliquoted into PCR tubes into which a small amount of yeast cells were transferred using the end of a 200 μL micropipet tip. A standard hot-start thermocycler program with extension at 68°C was adjusted for the Tm of the primer set and size of the amplification product.

### Yeast Transformation

Transformations of *med15Δ* strains were conducted using the Frozen-EZ Yeast Transformation II kit (Zymo Research) with modifications. 10 mL of early log phase cells were pelleted and washed with 2 mL EZ 1 solution, repelleted and resuspended in 1 mL EZ 2 solution. Aliquots were frozen at -80°C. 0.2-1 μg DNA and 100 μL EZ 3 solution were added to 10 μL competent cells for each transformation. Transformations were plated on selective media following incubation at 30°C for 45 minutes. Transformations into all other strains were conducted using a standard lithium acetate transformation protocol (30, 31).

### Small Scale Fermentation Reactions

White grape juice concentrate (Global Vinters Inc., Canada) was supplied at ∼68°Bx (1°Bx is equal to 1 g/100 mL) and was diluted to 20-21°Bx with distilled deionized water as measured using a triple scale hydrometer and confirmed with a refractometer (Milwaukee). Grape juice media was supplemented with 200 mg/L histidine, 300 mg/L leucine, 300 mg/L lysine, and 200 mg/L uracil for use in growth and fermentation experiments.

10 mL molded sterile clear glass vials (American Clinical Supplies Inc.) were used in fermentation experiments. 5 mL of prepared grape must was added to each vial which was then closed tightly with a rubber cap. A 25 G x 7/8 inch (0.5 mm x 22 mm) hypodermic needle was inserted into the cap to allow CO_2_ release from the system and to restrict evaporation (8). Prior to inoculating the fermentation vials, cells were grown in SC-Leu media for 2.5 days to reach saturation and then diluted 1:20 into fresh SC-Leu media. 1.25x10^7^ cells of saturated culture was added to 5 mL prepared white grape must (or other media as indicated in the figure legends) to a final cell concentration of 2.5x10^6^ cells/mL. SC-Leu media was added as needed to standardize the media added to each vial. Uninoculated vials with the same volume of grape must were used to control for contamination. Fermentation vials were incubated at 25 ± 1°C or as indicated. The temperature was maintained at 25°C despite fluctuating room temperatures using a heated shaker in a 4°C cold room. Shaker speed was maintained at 165 rpm.

### Fermentation Assays

Progression through the fermentation process was determined by once-daily weight loss measurements caused by CO_2_ release. Cumulative CO_2_ weight loss was calculated by subtracting daily measurements from the initial weight (day 0). To examine the impact of each *MED15* plasmid, fermentation (weight loss) was evaluated simultaneously for multiple different transformants (biological replicates). Time to weight loss benchmarks were determined per transformant. The total amount of CO_2_ weight loss accumulated by day 7 for *MED15* strains or day 8 for *med15Δ* strains was set as 100% weight loss. Values equal to 10%, 35%, 50%, and 80% of the 100% value were plotted as horizontal lines on top of fermentation curves. The time, in hours, that corresponded to the intersection between the horizontal lines and fermentation curves was determined by eye.

### RNA Extraction

RNA was extracted from 10 mL yeast cultures harvested in log phase (2x10^7^ cells/mL). Cells were pelleted by centrifugation and resuspended in 1 mL cold water. Yeast were repelleted by centrifuging for 10 seconds at highest setting of tabletop centrifuge. The supernatant was aspirated, and the pellet was frozen in dry ice for 5 minutes before being stored at -80°C prior to RNA extraction.

RNA was extracted using a hot acid phenol protocol (32). Frozen cell pellets were resuspended in 400 µL TES solution (10 mM Tris, 10 mM EDTA, 0.5% SDS). An equal volume of acid phenol was added to each tube and vortexed for 10 seconds. Tubes were incubated in a 65°C water bath for 1 hour with vortexing every 10-12 minutes. Phases were separated by microcentrifugation in the cold for 5 minutes. The aqueous layer was extracted into a new tube avoiding the DNA-enriched interface. A second round of acid phenol extraction was conducted followed by a chloroform extraction. RNA was precipitated with 300 µL of 4 M LiCl in dry ice for 20 minutes followed by 5 minutes of high-speed centrifugation at 4°C. The pellet was washed with 500 µL ice cold 70% ethanol and dried for 10 minutes at 37°C. RNA pellets were resuspended in DEPC treated water and stored at -20°C. Contaminating genomic DNA was removed using the DNase Max Kit (Qiagen). 10 µL of a 1x Master Mix (5 µL 10X Buffer, 5 µL water, and 0.5 µL DNase I) was added to 40 µL RNA. Following a 30-minute incubation at 37°C, DNase was removed by addition of 5 µL of the DNase removal resin and a10 minute RT incubation with periodic agitation. The resin was pelleted by centrifugation for one minute at the highest setting of a microcentrifuge. The supernatant containing DNA-free RNA was transferred to a new tube and the concentration determined using a NanoDrop spectrophotometer (Thermo Scientific). RNA quality was visualized on formaldehyde gels (1% agarose, 10% formaldehyde solution, 1x MOPS buffer).

### cDNA Preparation and qRT-PCR

cDNA was prepared using Superscript III First-Strand Synthesis System (Invitrogen). Transcripts were amplified with random hexamer (Invitrogen) or anchored oligo-dT20 (Integrated DNA Technologies) primers. For anchored oligo-dT20 primers 1 μg RNA, 50 ng primer, and 0.01 μmol dNTPs were mixed in a final volume of 10 μL. Tubes were incubated for 5 minutes at 65°C and then 2 minutes on ice. 10 μL cDNA Master Mix (2x RT buffer, 10 mM MgCl_2_, 0.02 M DTT, 40 U RNase OUT, 200 U Superscript III Reverse Transcriptase) or mock Master Mix lacking reverse transcriptase was added to each tube and incubated for 60 minutes at 50°C and 5 minutes at 85°C. To degrade RNA, 2 U RNase H was added to each tube and incubated for 20 minutes at 37°C. For random hexamer primers tubes were incubated 10 minutes at 25°C and then 50 minutes at 50°C.

Transcript abundance was quantified for each sample relative to a normalization transcript using the PerfeCTa SYBR Green FastMix (Quantabio) in 96-well PCR plates (Hard-Shell 480 PCR Plates, Bio-Rad) measured in a LightCycler 480 (Roche). Individual 10 µL reactions consisted of 5 µL FastMix, 1 µL target-specific primer pairs (0.5 µL 10 µM forward primer and 0.5 µL 10 µM reverse primer), 3 µL DEPC treated water, and 1 µL cDNA, mock cDNA, or water. Plates were sealed with optically clear film (PlateSeal). A standard SYBR green PCR program was used with modifications: 95°C for 5 minutes, 45 cycles of 95°C for 10 seconds, 55°C for 10 seconds, and 72°C for 20 seconds with a single fluorescence acquisition. A melting curve was conducted directly following the PCR program to confirm the presence of individual species amplified in each well. The melting curve program was 95°C for 5 seconds, 65°C for 1 minute, and ramp up to 97°C by 0.11°C/s with continuous fluorescence acquisition. *ALG9* was used as a normalization transcript. All samples were measured with three technical replicates. Mock cDNA (prepared without the addition of reverse transcriptase) and water were used as negative controls.

The relative abundance of target transcripts was determined by calculating the crossing point (CP) value for each reaction which reflects the number of cycles of amplification required for the SYBR green signal to exceed a threshold value set to eliminate background noise. CP values were calculated using the Second Derivative Maximum method implemented in the LightCycler 480 Software (Roche). The average CP for technical replicates amplifying the target transcript with a specific RNA sample was normalized to the average CP for technical replicates amplifying the normalization transcript. This ratio was then compared across samples as a depiction of relative abundance of the target transcript in each sample.

### RNA Sequencing

Individual fermentation reactions in 5 mL supplemented white grape juice were prepared as previously described for two biological replicates each of the *URA3* marked *med15Δ* strain and integrants of the LAB, WY7, WY15, and WY23 *MED15* alleles. Weight loss was monitored, and yeast were pelleted at the 35% weight loss point (∼37 hours for *MED15* strains, and 48 hours for *med15Δ* strains). RNA was extracted as described above. RNA sample quality was assessed using an Experion automated electrophoresis system with a HighSens chip using a standard protocol (Bio-Rad). cDNA libraries were prepared using the Illumina Stranded mRNA Prep protocol (Illumina). A total of 150-200 ng of total RNA per sample and 13 cycles of amplification were used. The Illumina Dual Index Adapters set A (Integrated DNA Technologies) was ligated to the cDNA.

cDNA library quality control and sequencing were performed by the Iowa Institute for Human Genetics (IIHG) Genomics Division. cDNA libraries were quantified using fluorometry with PicoGreen. The integrity of the cDNA libraries was determined using an Agilent BioAnalyzer DNA chip. A KAPA qPCR was used to get precise quantification prior to loading samples for sequencing. The pooled libraries were run with the 2x150 bp protocol on one lane of an SP flow cell in an Illumina NovaSeq.

Sequencing data was processed using a manually curated pipeline of programs implemented in the IIHG instance of the Galaxy platform, a web-based computational workbench (33). Adapter sequences were trimmed from reads using TrimGalore! (Galaxy tool version 0.6.3). The adapter sequence was auto detected. Reads shorter than 50 bp after trimming were discarded. Any reads orphaned at this point were also discarded to maintain sets of fully paired reads. Paired-end reads were then mapped to the S288C reference genome (sacCer3) using Bowtie2 (Galaxy tool version 2.3.4.2) (34). Default parameters for paired-end reads were used, the maximum paired-end length was set to 5000 and dovetailing allowed to maximize the number of mapped reads. Counts were quantified using featureCounts (Galaxy tool version 1.6.3) (35) to tally the number of reads mapping to annotated “genes” in the S288C reference general feature format file (version R64-3-1) obtained from downloads.yeastgenome.org/latest/saccharomyces_cerevisiae.gff.gz. Count normalization and differentially expression analysis was done using DESeq2 (Galaxy tool version 2.11.40.2) (36) using default parameters with multiple testing correction using Benjamini-Hochberg method with a false discovery rate of 5%.

Volcano plots of differentially expressed genes were prepared using VolcaNoseR (37). Venn diagrams of overlapping differentially expressed genes were prepared using BioVenn (38). GO-enrichment analyses were done using Yeastmine (39) and Enricher (40–42). Metabolic pathway enrichment analyses were done using BioCyc (43).

### Gene Set Enrichment Analysis

Pathways enriched in differentially expressed genes, in comparisons of *MED15*_WY_ alleles with *MED15*_LAB_, were identified using fgsea (44). Wald statistics for differential expression from the DESeq2 analyses were used to determine enrichment among 1,234 yeast pathways compiled in the ConsensusPathDB (45). These included pathways from BioCyc, KEGG, and Reactome among others. For each comparison, pathways were excluded from the analysis if there were five or fewer genes with Wald statistics within that pathway. Pathways with adjusted p-values less than 0.01 were considered significantly enriched. Venn diagrams of overlapping enriched pathways were created using VennDiagram (46). Heatmaps for gene sets that correspond to specific pathways were created using pheatmap (47) for log transformed normalized counts.

### Primers

All primers were synthesized by Integrated DNA Technologies and are listed in Tables 3 and 4. Primer pair efficiency was measured by qRT-PCR analysis of serial dilution of control RNA.

**Table 3.**
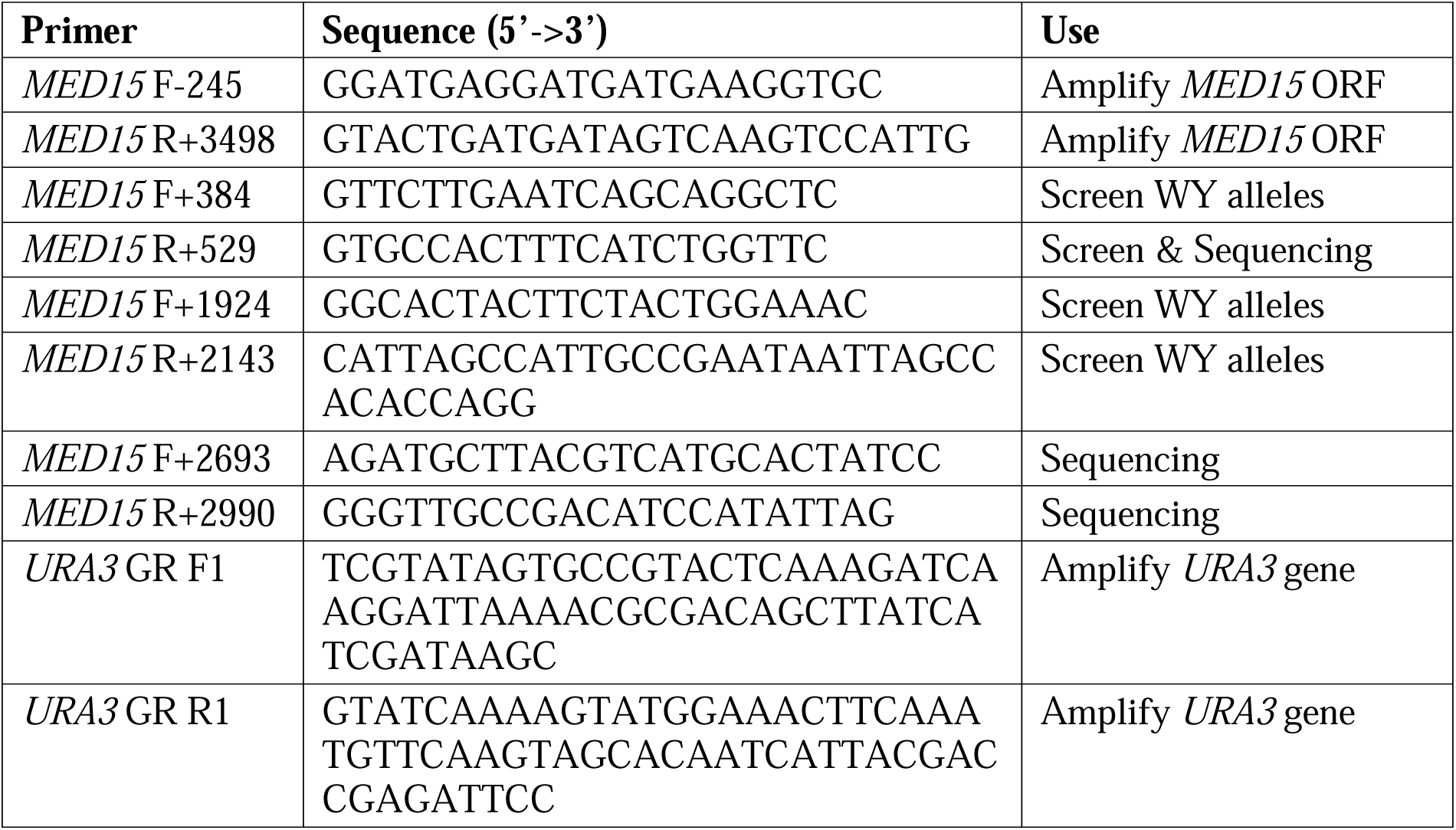
Primers.

**Table 4.**
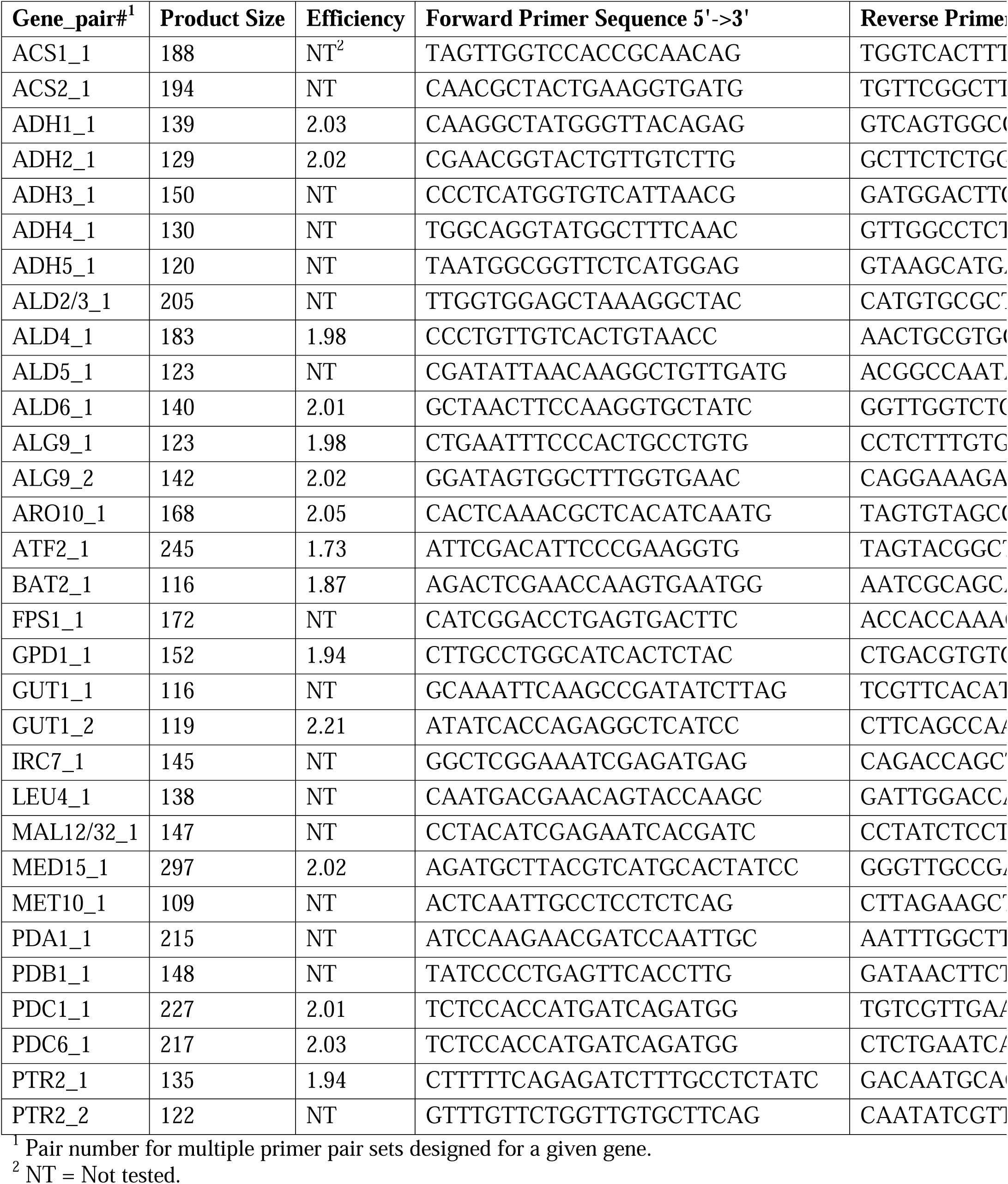
qRT-PCR Primers.

## Results

### Fermentation Gene Expression is Affected in the *med15Δ* Strain

We previously observed that a *med15Δ* lab strain is defective in the fermentation of grape juice (1). We hypothesized that this fermentation defect is the consequence of changes in the expression of metabolic genes involved in the production or consumption of ethanol. To determine the role of Med15 in the regulation of relevant fermentation genes, we first analyzed microarray datasets (48–50) to identify genes with previously described roles in ethanol metabolism as well as in wine and beer fermentation (51–53) and evaluated their expression. Many genes with roles in ethanol metabolism and fermentation were regulated by *MED15* including three (*PDC1*, *ACS1*, and *ADH2*) that had not previously been recognized as *MED15* regulated genes in published microarray datasets. Both positive regulation by Med15 (lower expression in the deletion strain) and negative regulation by Med15 (higher expression in the deletion strain) was observed. The targeted gene-by-gene qRT-PCR data we report here (Fig. 1B) is expected to be more robust than global microarray datasets in the literature (48–50) that we present for comparison (Table S1, columns 3-5).

**Figure 1.**
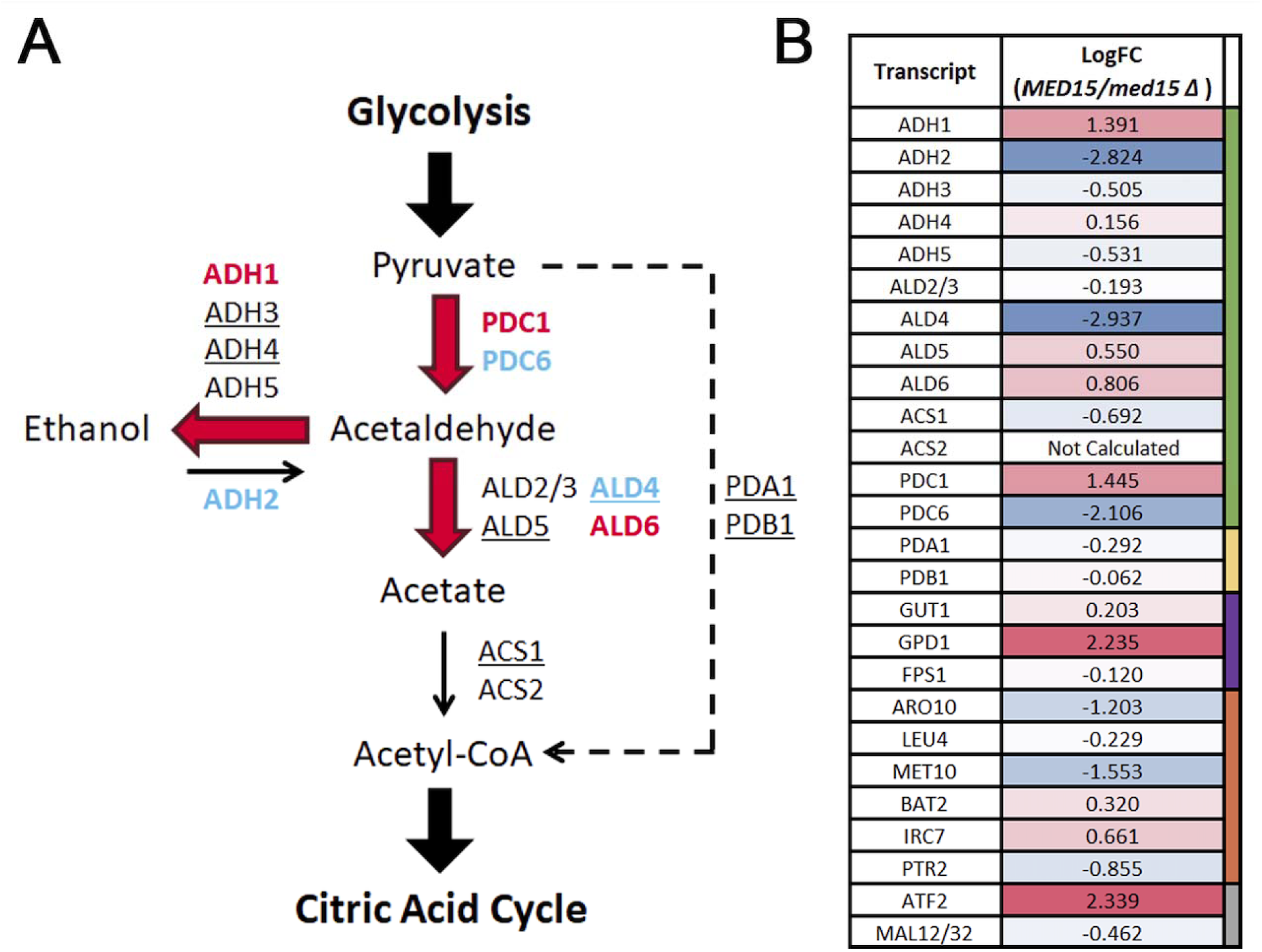
Influence of Med15 on expression of alcoholic fermentation pathway genes. **(A)** Summary of predicted impact of *med15Δ* on ethanol metabolism. Tested metabolic genes labeled according to changes in expression in *med15Δ* relative to expression in WT yeast. Gene names in red are upregulated (activated) by *MED15*, gene names in blue are downregulated (repressed) by *MED15*, and gene names in black not regulated by *MED15*. Underlined genes are mitochondrial. **(B)** Gene expression normalized to *ALG9* in *MED15* (set to 1) relative to *med15Δ* for 3-6 technical replicates. Relative expression values for genes with shades or red representing degrees of upregulation (activated) by *MED15*, and shades of blue representing downregulation (repressed) by *MED15*.

The Med15-regulated gene list includes a subset of genes involved in ethanol metabolism, specifically the alcohol dehydrogenases *ADH1* and *ADH2*, and the aldehyde dehydrogenases *ALD4* and *ALD6*. Other ADH and ALD genes were not significantly affected by Med15. In the *med15*Δ strain, expression of *ADH1*—which encodes an enzyme that converts acetaldehyde to ethanol—was reduced, while *ADH2*, which catalyzes the reverse reaction (ethanol to acetaldehyde), was significantly upregulated (Fig. 1). This suggests that Med15 normally promotes *ADH1* expression and represses *ADH2*. Similarly, the ALD genes, which convert acetaldehyde to acetate, showed opposing regulation. *ALD4* expression is increased, while *ALD6* expression is decreased in the *med15*Δ strain (Fig. 1). These results indicate that Med15 typically represses the mitochondrial, glucose-repressed *ALD4* and induces the cytoplasmic *ALD6*. A similar pattern is observed with the pyruvate decarboxylases. *PDC1*, the major isoform responsible for converting pyruvate to acetaldehyde during fermentation, is downregulated in the *med15*Δ strain, while *PDC6*, a minor isoform typically expressed under stress conditions, is upregulated. This suggests that Med15 promotes *PDC1* and represses *PDC6*, reinforcing a fermentative metabolic profile. Together, these effects suggest that Med15 plays a central role in fermentation by regulating key enzymes that drive ethanol production, ethanol degradation and stress-related metabolic shifts.

### The *MED15* Transcriptome is Highly Specialized during Alcoholic Fermentation

To determine the global impact of *med15Δ* on gene expression during alcoholic fermentation rather than in rich media (YPD, 2% glucose) as in the datasets discussed above, an RNA Sequencing experiment was conducted for cells grown in white grape juice (WGJ; 20% glucose) at the 35% weight loss fermentation benchmark (Fig. 2A) (1, 8). Note that weight loss benchmarks rather than time was used to coordinate sample collection in this experiment because the *med15Δ* strain grows and ferments much slower than strains with an intact *MED15* gene. This global analysis was expected to identify all affected genes including any that we had not anticipated, and the fermentation conditions were expected to have a substantial impact on gene expression.

**Figure 2.**
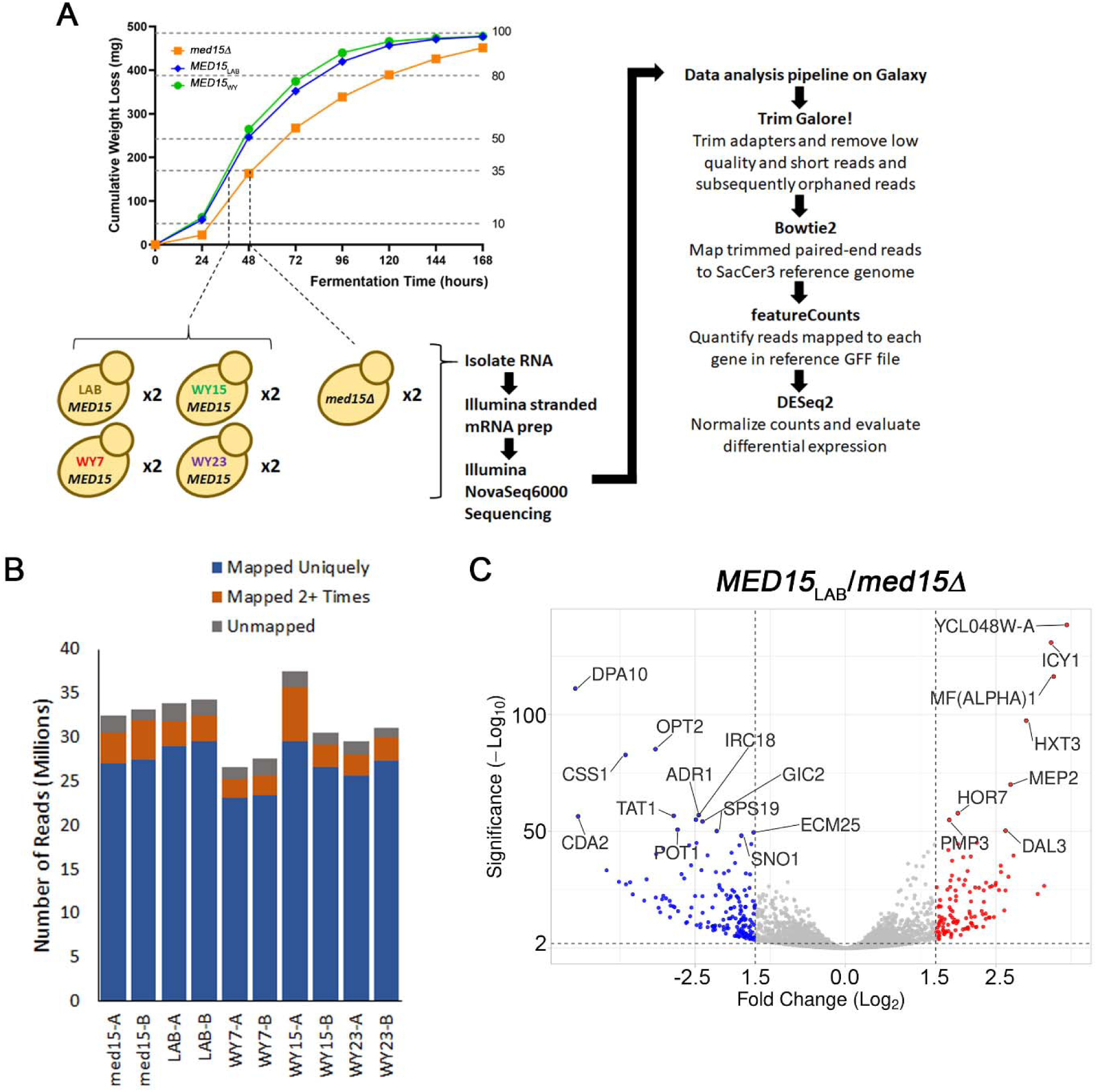
WGJ RNA Sequencing experimental setup and summary results. **(A)** RNA sequencing experimental overview. **B)** Summary results from Bowtie2 mapping for each sample. **C)** Volcano plot of differentially regulated genes between LAB and *med15Δ* samples.

In this analysis we found 615 genes that were differentially regulated in a comparison of WT to the *med15Δ* strain (log2FC ≥ 1 or ≤ -1 and adjusted p ≤ 0.001). The most highly differentially expressed genes (n=282, log2FC ≥ 1.5 or ≤ -1.5, and adjusted p ≤ 0.0001) are both positively and negatively regulated by Med15 (Fig. 2C). Surprisingly only about 20% of differentially regulated genes overlapped with genes differentially regulated by *MED15* in the non-fermentation (YPD) environment (Fig. 3A). Condition-nonspecific functions regulated by *MED15* were identified by gene ontology (GO) enrichment analysis of genes identified in both conditions. Enriched terms included: amino acid biosynthesis, carbohydrate metabolism, cell wall organization, and transport (Fig. 3B).

**Figure 3.**
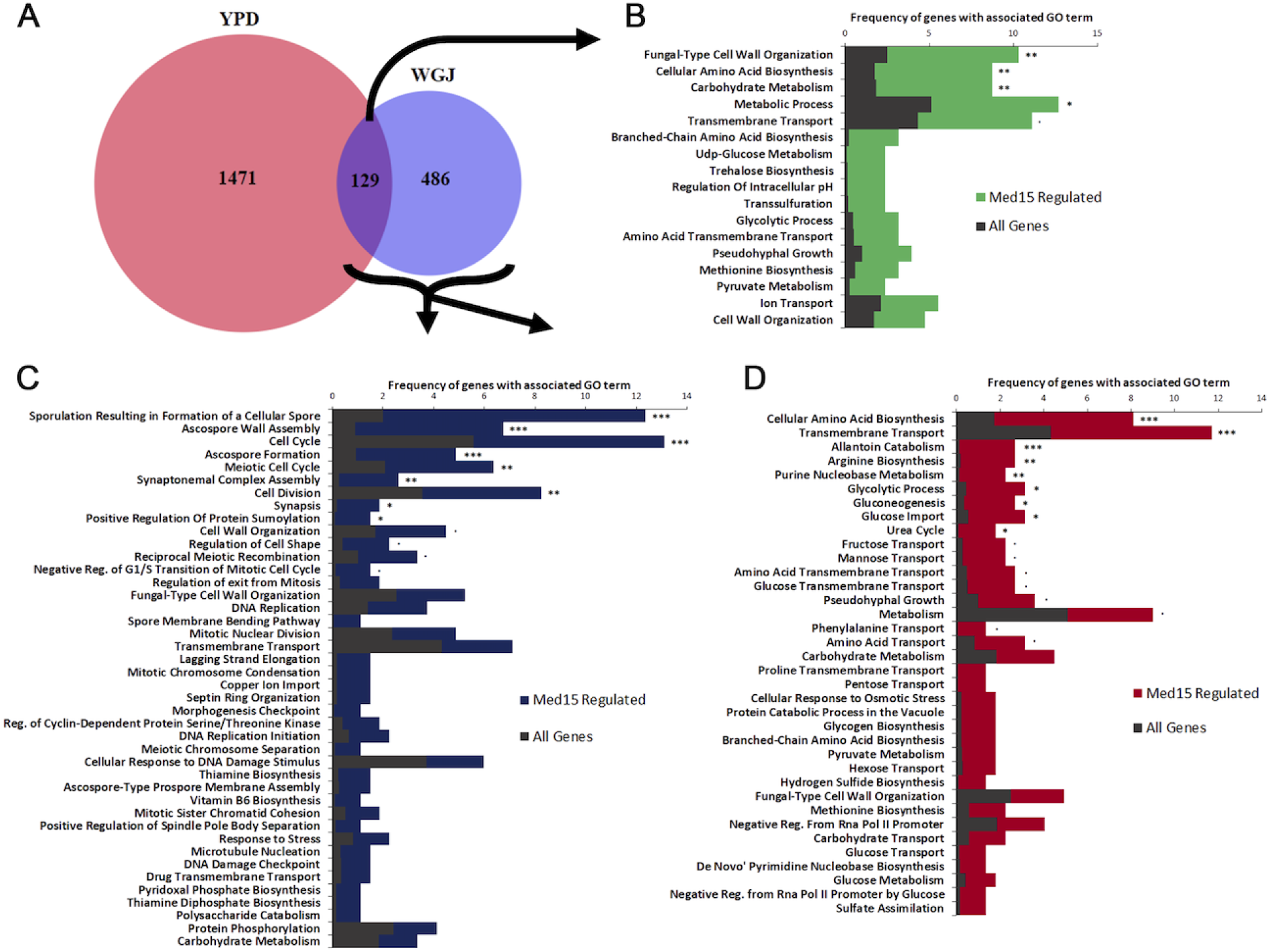
Condition specific and general metabolic pathways regulated by *MED15*. **(A)** Overlap between differentially regulated genes in the *med15Δ* strain: from Hu *et al.* (2007) microarray data (48) for log phase growth in YPD (YPD) and from RNA sequencing data at the 35% weight loss benchmark in WGJ fermentation (WGJ). *MED15* regulated genes were defined as log2 fold change values of >1 or <-1 for *med15Δ* vs WT. **B)** GO enrichment for genes regulated by *MED15* in both datasets (from overlap in Venn diagram). GO enrichment analysis was conducted with YeastMine in SGD (39). **(C)** WGJ fermentation GO Enrichment for genes negatively regulated by *MED15*. GO enrichment for genes more highly expressed in *med15Δ* in the WGJ fermentation environment. GO enrichment analysis done using YeastMine in SGD (39). **(D)** WGJ fermentation GO Enrichment for genes positively regulated by *MED15.* GO enrichment for genes more highly expressed in *MED15* in the WGJ fermentation environment. GO enrichment analysis done using YeastMine in SGD (39). For (B), (C), (D), listings only include nominal enrichment P values of <0.05. Adjusted P value intervals include • < 0.1; * < 0.05; ** < 0.01; *** <0.001.

Genes unique to the white grape juice (WGJ) environment and negatively regulated (significantly lower expression in *MED15* vs *med15Δ*) included: sporulation, cell cycle (DNA replication, cell division, checkpoints, and exit from mitosis), cell wall organization, response to stress, and transport (copper ions, iron ions, and drugs) (Fig. 3C). Genes positively regulated by Med15 in WGJ were enriched in carbohydrate transport (pentose, hexose, glucose, fructose, and mannose), glycolysis, amino acid transport, and amino acid biosynthesis (Fig. 3D). Seven *HXT* genes were found to be *MED15* regulated, including the activation of high-affinity (*HXT4*, *HXT6*, and *HXT7*) and low-affinity (*HXT1* and *HXT3*) glucose transporters, and repression of *HXT10* and *HXT14*.

To achieve more nuanced insight into the *MED15*-dependent metabolic profile during fermentation, we analyzed the distribution of *MED15* regulated genes across all yeast metabolic pathways using the BioCyc database (43) with the entire WT vs *med15Δ* fermentation dataset as input (including positively regulated, negatively regulated, and genes not regulated by *MED15*). In addition to glycolytic and ethanol metabolic genes, the arginine biosynthetic pathway was specifically enriched during WGJ fermentation (*ARG1, ARG3, ARG4, ARG5* and *ARG6*) with all regulated genes expressed more highly in the *MED15* strain (Fig. 4C). A similar enrichment was seen for genes involved in the methionine biosynthetic and allantoin degradation pathways (Fig. 4D, E). Allantoin catabolism was one of the most significant hits for BioCyc pathways regulated by *MED15* during fermentation. All allantoin catabolism enzymes (*DAL1*, *DAL2*, *DAL3*, and *DUR1,2*) were significantly upregulated by *MED1*5.

**Figure 4.**
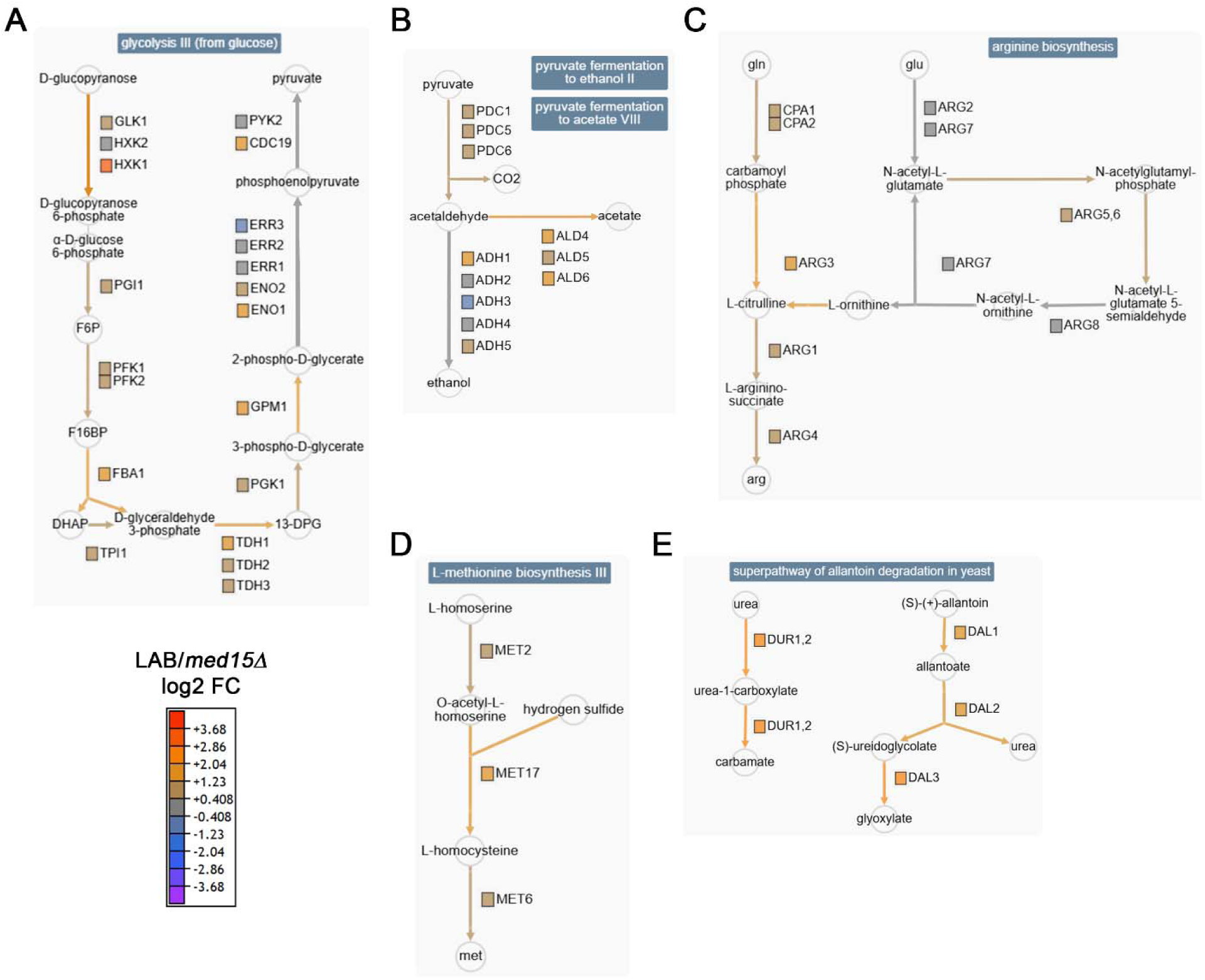
Metabolic pathways regulated by Med15. **(A)** glycolysis; **(B)** pyruvate fermentation; **(C)** arginine biosynthesis; **(D)** methionine metabolism; and **(E)** allantoin metabolism. Pathways are generated using BioCyc (43). The color of the edges (arrows) are approximations of the sum of the effects on the individual steps (enzymes) in the pathway. See key for the relative Med15 dependence of each enzyme in the pathway.

Because published reports show that *MED15* affects silencing at telomeres during normal growth (54, 55), subtelomerically localized genes were evaluated in the alcoholic fermentation dataset. We evaluated the distal region of yeast telomeres which contains genes belonging to a small number of gene families including *MAL*, *MEL*, *PAU*, *COS*, and *FLO* genes (Fig. 5) although the S288C lab strain used here lacks *MEL* genes (56) and only has two partially functional *MAL* gene regulons (57, 58). Most *PAU* genes (12/21) and *COS* genes (7/11) and all *FLO* genes were significantly differentially regulated by *MED15* under fermentation conditions. Virtually all *FLO*, *PAU*, and *COS* genes (except for *PAU9*) which were significantly differentially regulated between the WT and *med15Δ* strains were more highly expressed in the *med15Δ* strain. This pattern suggests that during fermentation *MED15* promotes or helps maintain subtelomeric gene silencing. This is in contrast to the expression pattern of subtelomeric genes during log-phase growth in YPD during which only a few of these genes are regulated by *MED15* with an average fold change across genes in each family showing higher expression in the WT strain (48–50).

**Figure 5.**
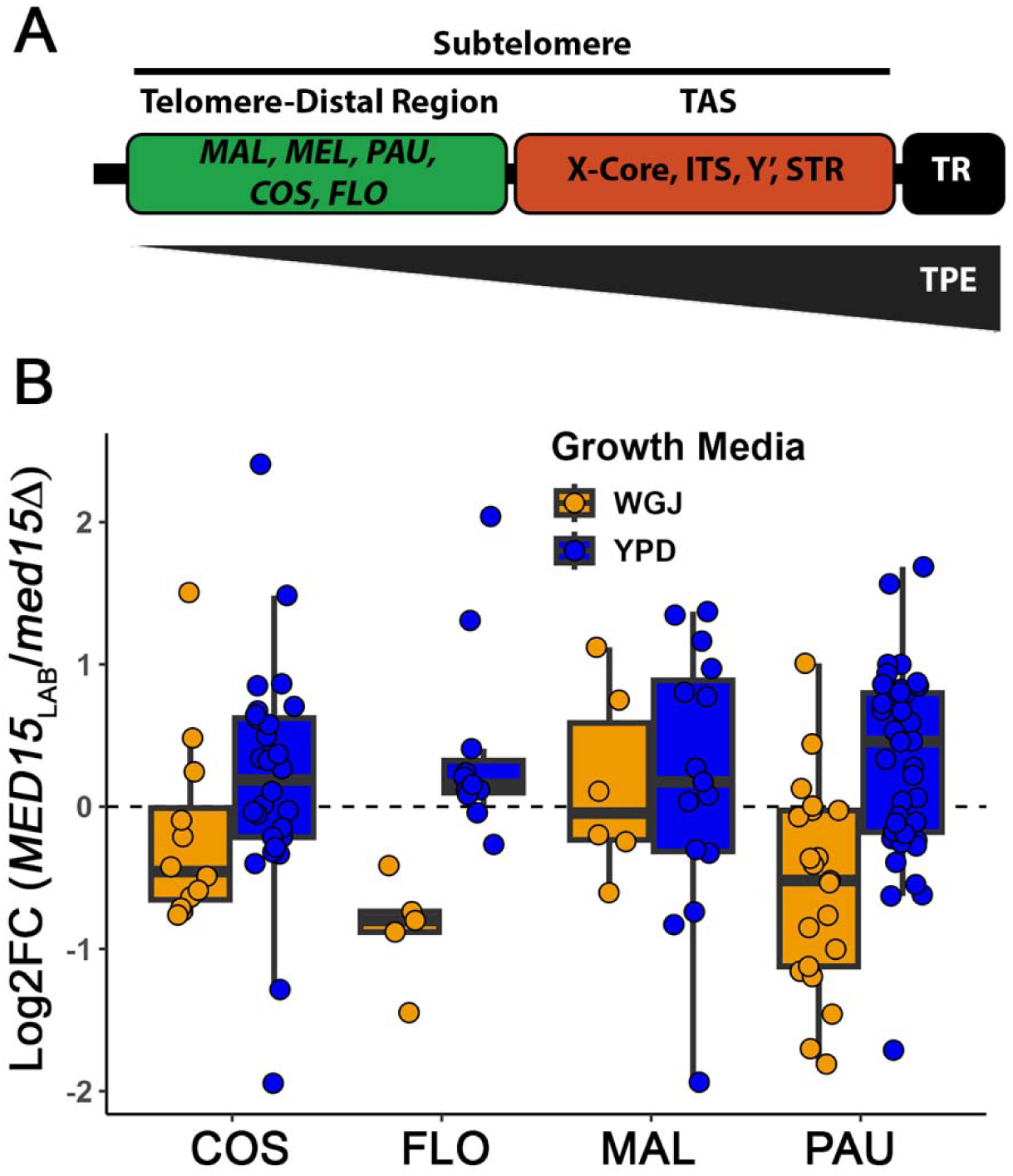
*MED15* regulates silencing of subtelomeric genes during WGJ fermentation. **(A)** Cartoon depiction of *S. cerevisiae* telomeres and subtelomeric features. Abbreviations: Telomeric Repeats (TR), Telomere Associated Sequence (TAS), Interspersed Telomeric Repeats (ITS), SubTelomeric Repeat sequences (STR) and Telomere Position Effect (TPE). **(B)** Differential expression of subtelomeric genes of each indicated class (COS, FLO, MAL, PAU) during WGJ fermentation and growth in YPD. Log2 fold change expression differences are shown for LAB/*med15Δ*. YPD data is from (48–50).

### *MED15*_WY_ - *MED15*_LAB_ Comparisons Reveal Subtle Effects on the Fermentation Transcriptome

To investigate the impact of wine yeast (WY) *MED15* alleles during alcoholic fermentation, we examined the impact of three different WY *MED15* alleles integrated into the LAB strain genome in RNA-Seq experiments at 35% weight loss (Fig. 2). Overall, the fermentation genes (labeled in Fig. 6) were not significantly affected. Only the *ALD* genes displayed absolute log2 FC values greater than 0.5 (Fig. 6 B, C and D) and only *ALD5* was significantly differentially regulated, and this was true only in WY23 (Fig. 6D). Thus, the expression of most fermentation genes was unchanged in WGJ environment regardless of *MED15* allele suggesting that the enhanced fermentation phenotypes in strains with *MED15*_WY_ alleles (1) is rooted in expression differences in other classes of genes.

**Figure 6.**
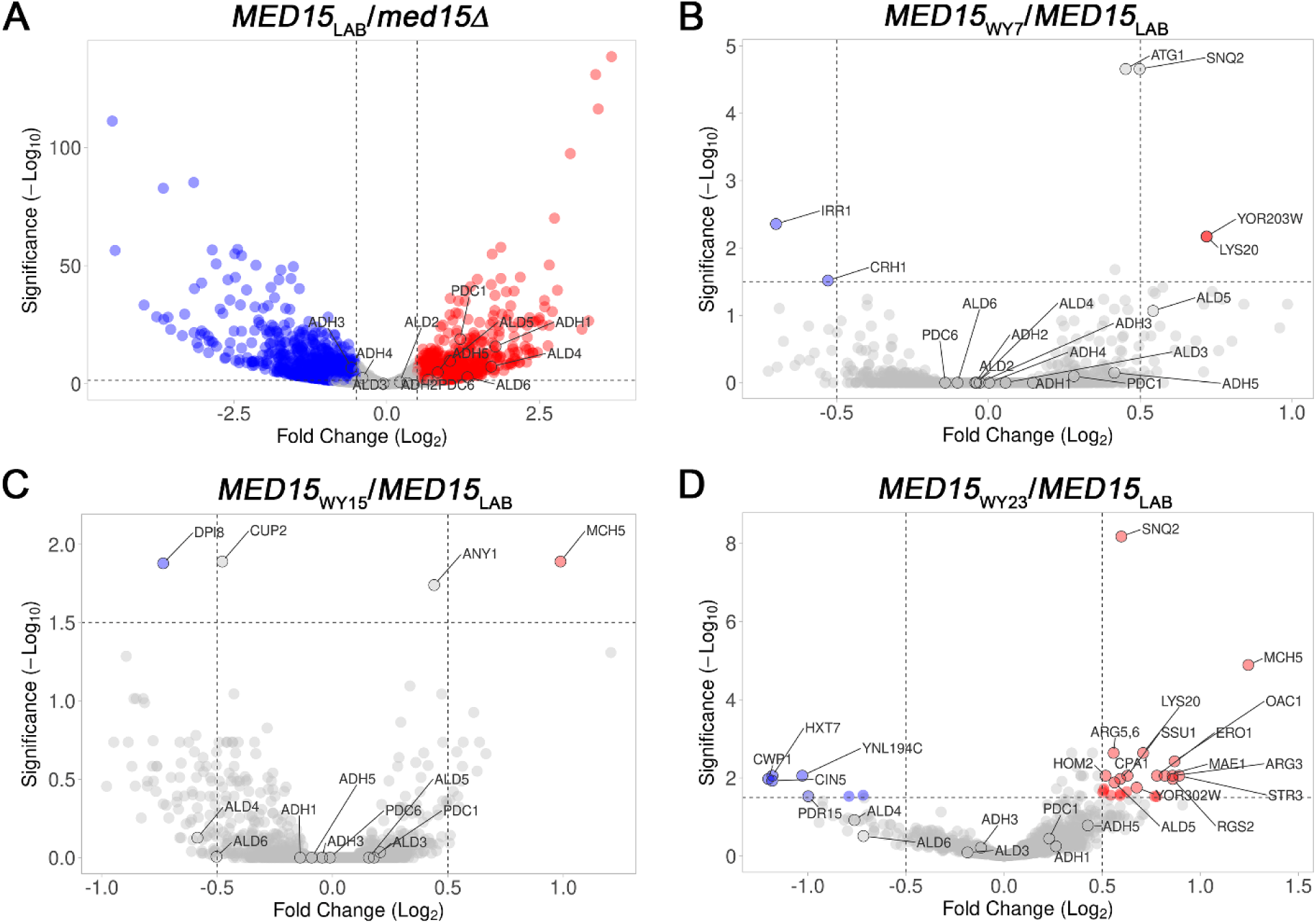
Fermentation metabolic gene expression in MED15_WY23_ vs MED15_LAB_. **(A-D)** Volcano plot of differentially regulated genes between strains expressing *MED15* from LAB and deletion (A), WY7 (B), WY15 (C) and WY23 (D) are shown. Highlighted in each plot are genes that exceed both fold change and P value thresholds represented by vertical and horizontal lines. Genes that are blue are negatively affected by the WY allele, and genes in red are positively affected by the WY allele. Each colored gene in the WY7, WY15 and WY23 plots is annotated. Additional annotations are for fermentation genes shown to be regulated (in YDP) as in Fig. 1.

To investigate the transcriptomic basis for improved alcoholic fermentation in strains with *MED15*_WY_ alleles (1), we used GO and Pathway enrichment analyses for the differentially regulated gene sets in comparisons of the transcriptomes of *MED15_WY_* and *MED15_LAB_* strains.

A comparison of strains with the WY23 *MED15* allele to strains with the LAB *MED15* allele in the WGJ environment revealed 31 differentially regulated genes (log2FC ≥ 0.5 or ≤ -0.5 and –log10 adjusted p ≥ 1.5) (Fig. 7, Table S2) and GO enrichment analysis pointed to an enrichment in amino acid metabolism and urea cycle genes suggesting an effect on nitrogen metabolism (Fig. 7). Specifically, the arginine biosynthetic pathway was upregulated due to the presence of *MED15*_WY23_*. MAE1* (malate dehydrogenase) expression was also significantly upregulated in the *MED15*_WY23_ strain alongside other enzymes in the gluconeogenesis metabolic pathway (Fig. 6D, Table S2).

**Figure 7.**
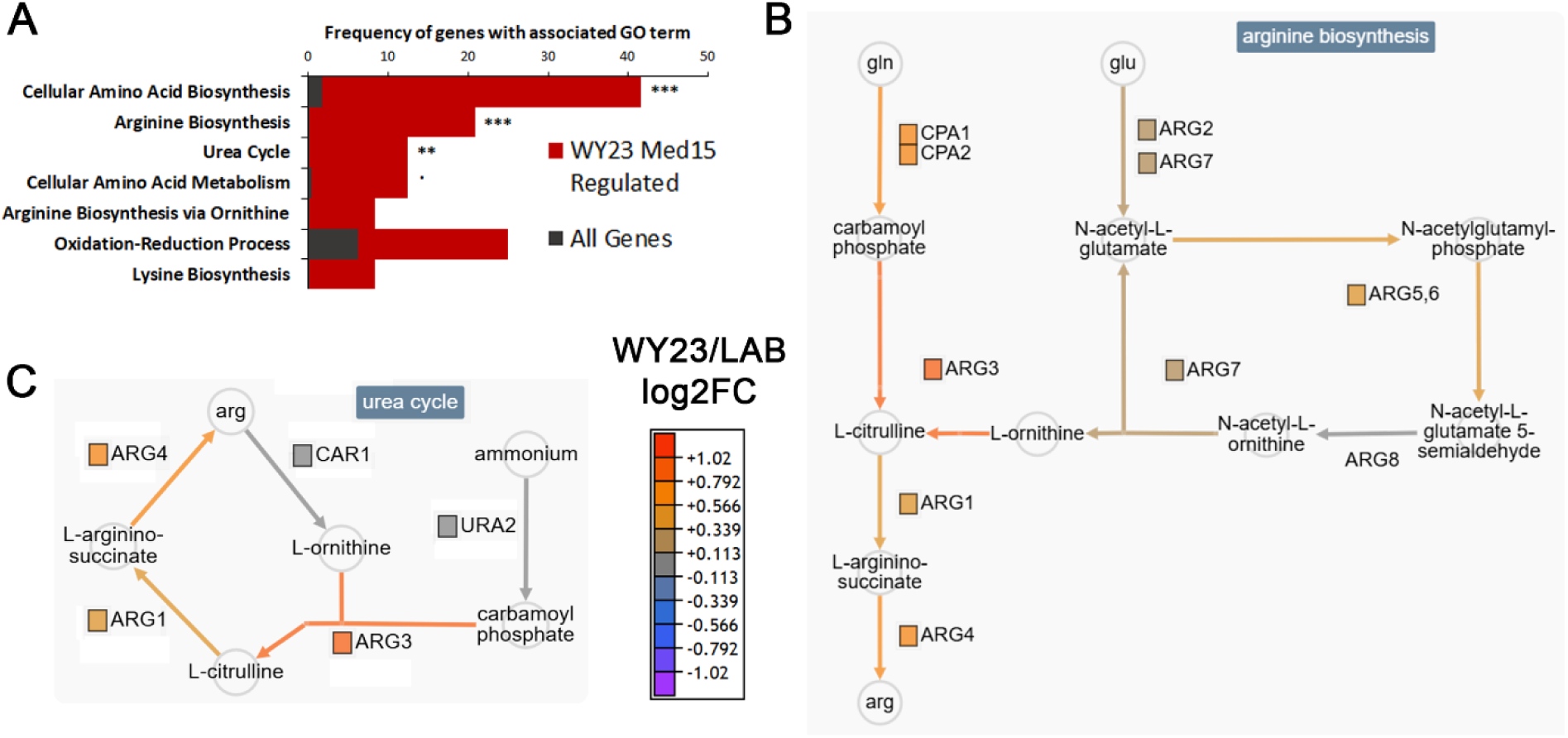
WY23 *MED15* differentially modulates the expression of metabolic genes during WGJ fermentation. **(A)** GO enrichment for genes more highly expressed in *MED15*_WY23_ in the WGJ fermentation environment. GO enrichment analysis conducted using YeastMine in SGD (39). Listings include nominal enrichment P values of <0.05. Adjusted P value intervals include • < 0.1; * < 0.05; ** < 0.01; *** <0.001. **(B-C)** Metabolic pathways for arginine biosynthesis (B) and urea cycle (C). Metabolic pathway analysis conducted with BioCyc (43). Edges of metabolic pathways are colored based on the activation of the pathway corresponding to the sum of the differential expression of the enzymes in that pathway.

Transcriptomes of strains with integrated *MED15*_WY7_ or *MED15*_WY15_ alleles were very similar to the *MED15*_LAB_ transcriptome at the fold-change and significance thresholds we set in Fig. 6. To achieve additional resolution, we combined GO term enrichment and gene set (pathway) enrichment analysis using slightly relaxed parameters. We also evaluated overlap among the enriched pathways in the different *MED15*_WY_ strains (compared to *MED15*_LAB_). The Venn diagram in Fig. 8 compares the pathways that are enriched in strains differing only in their *MED15* allele (*MED15*_WY23_, *MED15*_WY7_ and *MED15*_WY15_ and *med15*Δ) each compared to *MED15*_LAB_. We find for example, that the reduced expression of genes in the arginine pathway that we observed in the *med15* deletion strain relative to LAB (Fig. 4, Fig. 8 light blue) is mirrored by the elevated expression of many of the same genes in the comparison of *MED15*_WY23_ and *MED15*_LAB_ (Fig. 8, purple). This effect can also be seen in the heat map in Fig. 9.

**Figure 8.**
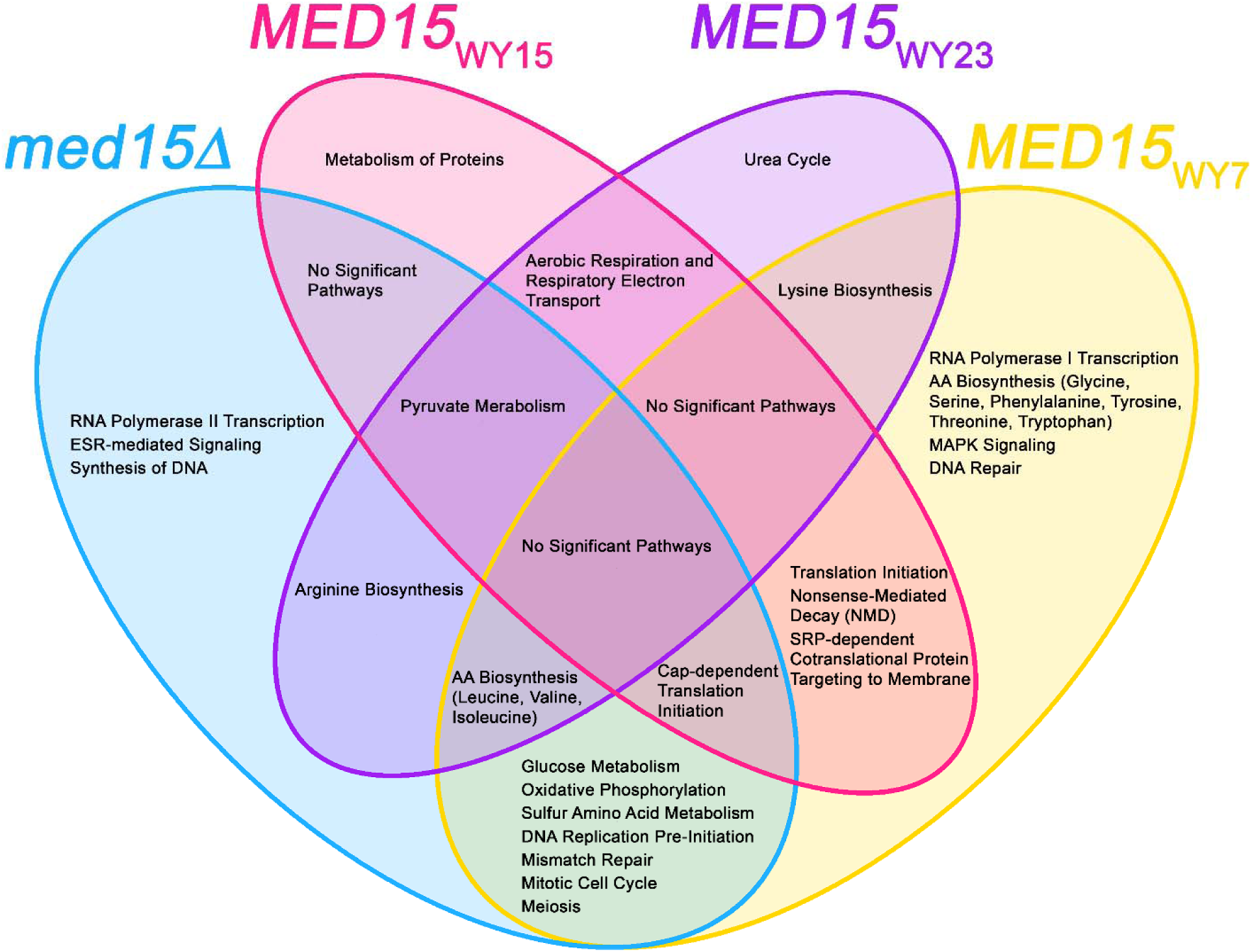
Venn Diagram depicting shared and unique pathways that are enriched in S288C based yeast strains each with different WY *MED15* allele. Genes regulated by *MED15* presence/absence in the lab strain are categorized in the blue segment of the diagram. Some of the same gene categories are affected in comparisons of strains with *MED15*_WY_ alleles compared to the *MED15*_LAB_ allele. For example, arginine biosynthesis genes are affected by the absence of *MED15* and by the *MED15*_WY23_ allele. Other affected gene categories are uniquely detected in comparisons of *MED15*_LAB_ strains to multiple *MED15*_WY_ alleles. For example, lysine biosynthesis is detected in *MED15*_WY23_ and *MED15*_WY7_ comparisons to LAB but is not among genes detected in a comparison between *MED15*_LAB_ and deletion strains. Finally, some gene categories (e.g., urea cycle genes) are unique to the *MED15*_WY23_ allele. This points to the existence of different gene expression patterns that may account for the enhanced fermentation characteristics exhibited by strains with *MED15*_WY_ alleles and to the WY strains themselves.

**Figure 9.**
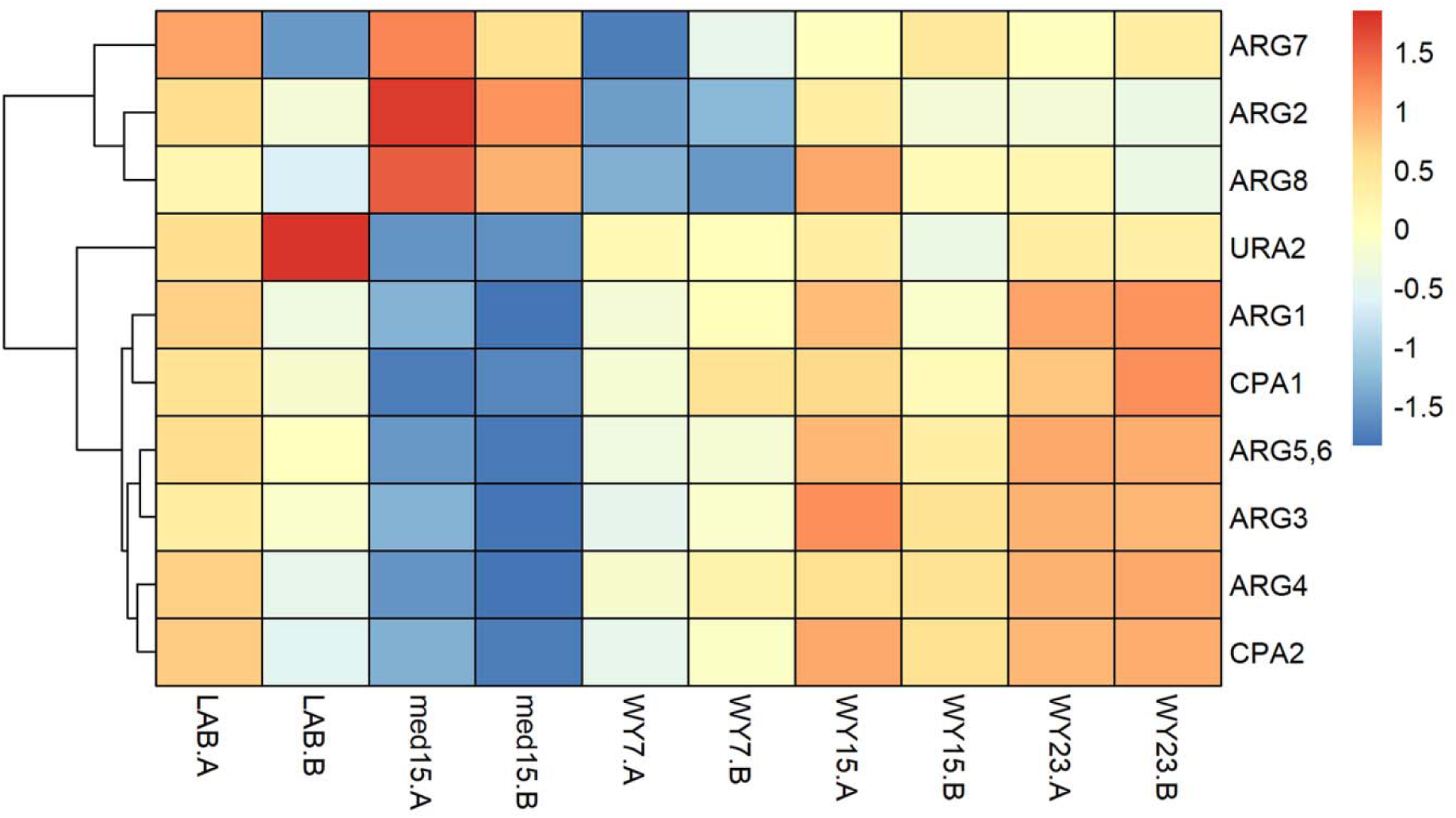
Heatmap for various pathways affected by *MED15* alleles during fermentation. These values are scaled per row from log2 transformed counts. Heat map representation of expression levels in strains with different MED15 alleles for various genes in the arginine biosynthesis pathway as defined by the YeastCyc database. Two biological replicates (A and B) of each strain type are shown. 7 of the 10 gene rows show relative downregulation in the *med15* mutant pointing to their normal activation by Med15, while *ARG7*, *ARG2* and *ARG8* are more highly expressed in the mutant, suggesting repression by Med15. Strikingly, these genes are underexpressed in strains with the *MED15*_WY7_ allele. Likewise, there is modest upregulation of *ARG1 – CPA2* (bottom 6 rows) relative to *ARG2*, *ARG7* and *ARG8* in strains with the *MED15*_WY23_ allele.

Similar overlap in pathways affected in the deletion (reduced) and in strains with *MED15*_WY7_ alleles (elevated) are seen for genes involved in DNA metabolism including meiosis, cell cycle, replication and repair (Fig. 8, intersection of the blue and yellow regions). Likewise, an effect on branched chain amino acid biosynthesis is seen in the deletion, *MED15*_WY7_, and *MED15*_WY23_ (Fig. 8, intersection of the blue, yellow and purple regions).

Some affected pathways in strains with WY *MED15* alleles are only minimally affected in the *med15Δ* mutant. For example, *MED15_WY7_* and *MED15_WY15_* have a shared set of affected translation functions ranging from translation initiation to nonsense-mediated decay and co-translational membrane targeting that are not prominent among genes affected in the *med15Δ* mutant. Likewise, the effect of *MED15_WY23_* and *MED15_WY7_* compared to *MED15*_LAB_ on lysine biosynthesis, or the effect of *MED15_WY23_* and *MED15_WY15_* compared to *MED15*_LAB_ on aerobic respiration and the urea cycle are not seen in the *med15Δ* mutant (Fig. 8)

### *MED15*-Dependent Arginine Regulation Affects Fermentation Efficiency

Given the pronounced effect of *MED15_WY23_* on arginine biosynthetic genes, we investigated the relevance of the arginine biosynthetic pathway in alcoholic fermentation. Fermentation experiments were conducted in SD medium with 20% glucose (plus amino acids required by the strains) and this media was supplemented with increasing amounts of arginine (Fig. 10 A, B). We found that arginine supplementation (to 800 μg/ml) reliably reduced the time to 50% weight loss by approximately 9% regardless of genotype (Fig. 10 B).

**Figure 10.**
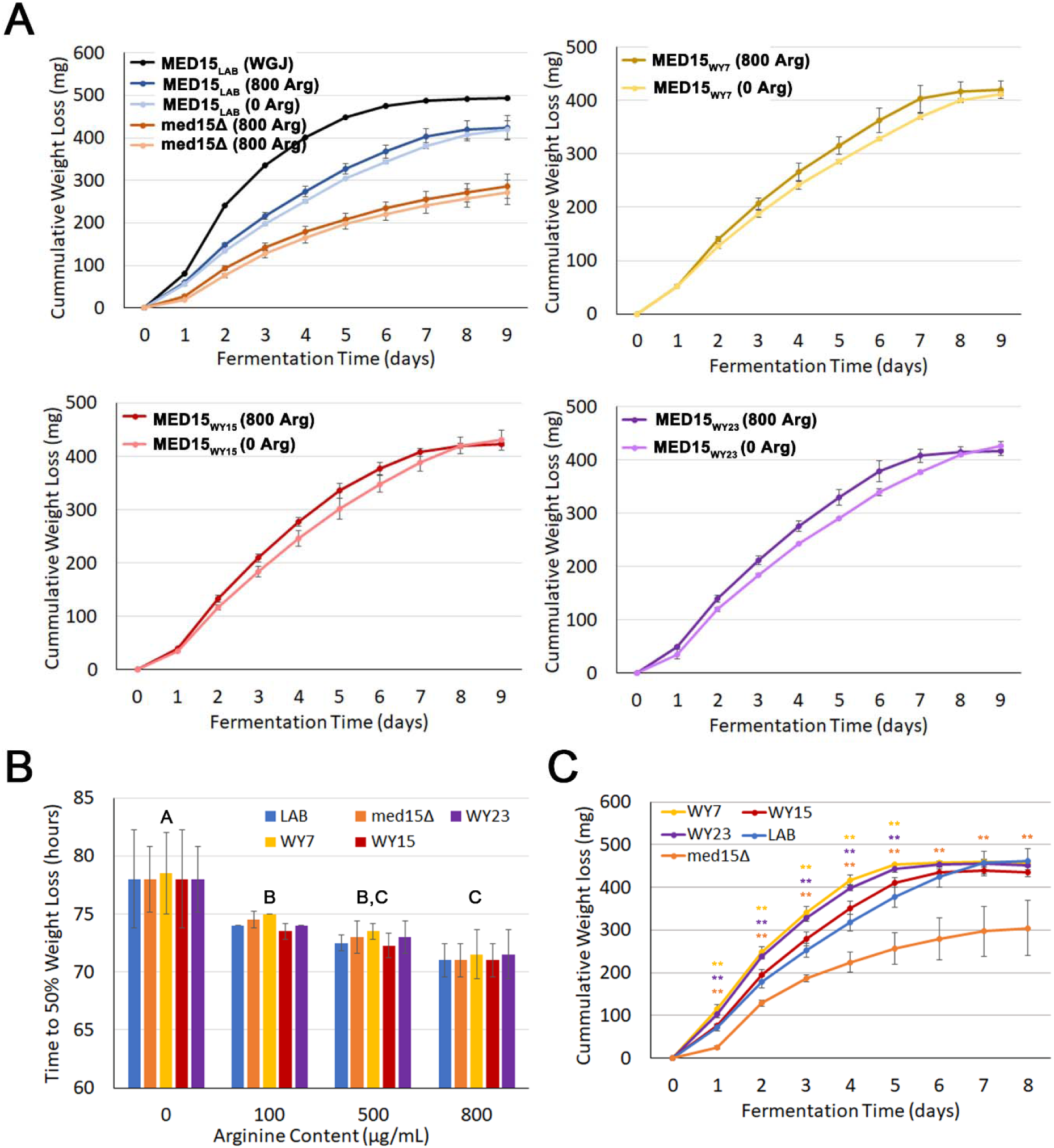
Improved fermentation among strains with MED15_WY_ alleles is evident in media lacking arginine. **(A)** Fermentation curves in SD media (lacking all but strain essential amino acids) with or without supplementation of arginine across strains with different *MED15* alleles. **(B)** Time to 50% complete fermentation in SD media with different levels of supplementation with arginine across strains with different *MED15* alleles. Significant differences were determined between arginine supplementation groups using ANOVA analysis with a Tukey post-hoc test, Group A – Groups B & C p<0.001, Group B – Group C p<0.01. **(C)** Fermentation curves in SC media without arginine (containing all other amino acids) across strains with different *MED15* alleles. The strains are the same as in (A) but abbreviated in (B) and (C) by their allele names for space. Significant differences between each strain and LAB determined using two-tailed t-tests, ** p<0.01

The impact of arginine limitation was evaluated by conducting fermentation in synthetic complete (SC with 20% glucose) media lacking arginine. We hypothesized that *MED15*_WY_ alleles might tolerate the absence of arginine better than strains with the *MED15*_LAB_ allele via altered expression of amino acid metabolic genes and better utilization of available nitrogen sources (1, 59). We found that strains with the *MED15*_WY7_ or *MED15*_WY23_ alleles clearly outperformed strains with the *MED15*_LAB_ allele in media lacking arginine, consistent with the observation that arginine biosynthetic genes are elevated in the *MED15*_WY23_ strain during fermentation (Fig. 10C).

## Discussion

Previously we found that *med15Δ* yeast strains were compromised in the efficiency of the fermentation process, while *MED15* alleles from wine yeast strains conferred a modest enhancement of fermentation when transplanted into a lab strain background (1). Here we examine gene expression in both YPD (qRT-PCR, Fig. 1) and during white grape juice-based alcoholic fermentation (RNA-Seq, Fig. 3-5) and show that *MED15* not only regulates relevant glycolytic, fermentation, and amino acid metabolic genes, but also genes that function in the DNA metabolism (DNA replication, repair) and translation (Fig. 8, *med15*).

### Transcriptional Patterns in YPD are Consistent with a role for *MED15* in Ethanol Production and Consumption

Previous analyses of the *med15Δ* transcriptomes in log-phase cells in YPD revealed that *MED15* regulates nearly a quarter of the genome (2, 49). qRT-PCR data (Fig. 1B) show that *MED15* plays a broad role in regulating levels of metabolic enzymes in many steps in the glycolytic and alcoholic fermentation pathways. The expression patterns suggest that *MED15* normally functions to promote ethanol production and consumption (Fig. 1). Expression data also indicated that *MED15* both positively and negatively regulates paralogous or redundant genes in multiple steps of these pathways during log-phase growth in YPD. For example, *MED15* positively regulates the major isoform *PDC1* and downregulates the minor isoform *PDC6*. Similarly, *MED15* downregulates the mitochondrial glucose-repressed *ALD4* and upregulates the cytoplasmic *ALD6*. The sum of *PDC* and *ALD* gene regulation would be expected to increase overall pyruvate decarboxylase and cytoplasmic aldehyde dehydrogenase activity.

*MED15* positively regulates *ADH1* and negative regulates *ADH2*. Adh2 preferentially converts ethanol back to acetate (17). Deletion of *ADH2* has been explored for strain engineering to increase the production of bioethanol (60). In quadruple mutants where the only alcohol dehydrogenase maintained is *ADH1*, yeast still produce wild type levels of ethanol (61). Therefore, considering the sum of *MED15*-dependent regulation of *ADH* genes, metabolism is expected to be skewed towards ethanol production. While winemakers and producers of bioethanol seek to engineer strains for ethanol production, our data suggests that natural variation in the general transcriptional regulator *MED15* has permitted evolution to accomplish that same task.

In addition to ethanol metabolism and fermentation genes, Med15 regulates glycerol and amino acid metabolic genes including *ATF2*, *GPD1*, *ARO10*, *MET10*, and to a lesser degree *IRC7* and *PTR2* during log-phase growth in YPD. Glycerol production is a desired property for wine and is beneficial for osmoregulation (6, 62). Strains have been engineered to produce less ethanol by the overexpression of *GPD1* to divert carbon to glycerol (53). Amino acid metabolism and peptide scavenging (*PTR2*) is critically important during fermentation in grape juice that is a nitrogen limiting growth medium (5, 6).

### Transcriptional Patterns under Alcoholic Fermentation Conditions

The *MED15* transcriptome is highly specialized during fermentation of WGJ. Only 20% of differentially regulated genes are identified in both YPD (2% glucose) and fermentation (20% glucose) datasets. Our RNA-Seq analysis confirmed positive regulation by Med15 of many glycolytic and alcoholic fermentation genes (Fig. 3, 4). The regulation of these metabolic genes is an obvious explanation for the fermentation deficiencies we previously described for the *med15Δ* strain (1). In addition to genes involved in the metabolism of glucose, *MED15* regulates a variety of disparate cellular activities during fermentation including cell wall homeostasis, cell cycle, sexual reproduction, carbohydrate transport, and amino acid biosynthesis (Fig. 3, 8). The regulation of these other cellular activities may also contribute to the fermentation phenotype of the *med15Δ* strain. For example, the yeast cell wall is known to change in composition depending on growth phase and the composition of the media (63, 64). Specifically, growth in nitrogen limiting media results in cell wall deformations potentially due to a reduced number of linkages between cell wall subunits (65).

Subtelomeric genes are silenced through heterochromatinization in a process referred to as the Telomere Position Effect (66). Our analysis of subtelomeric gene expression during WGJ fermentation indicates that *MED15* plays an enhanced role in telomeric silencing with 62% of analyzed subtelomeric genes having significantly lower expression in the WT strain compared to the *med15Δ* strain. Med15-dependent subtelomeric silencing has been previously reported (54, 55, 67), however, the effect on cells grown under laboratory conditions (YPD) is modest and in opposition to the effect during WGJ fermentation (Fig. 5B). This suggests that either *MED15*-dependent telomeric silencing is induced in grape juice fermentation but not YPD, or that *MED15*-independent silencing is present in YPD but not grape juice fermentation. During fermentation, reinforcement of TPE may serve to conserve cellular energy by limiting transcription of non-essential genes and stabilizing genome structure under stress.

The mechanism of enhanced subtelomeric gene silencing observed during fermentation may be related to the increased levels of H4K16Ac and reduced levels of Sir2 binding at telomeres in Mediator tail subunit deletions (67). Sir3 occupancy at telomeres is decreased in a *med15Δ* strain, which would correspond with a reduction in Sir2 binding (54, 55). Another potential mechanism by which Med15 could influence telomeric silencing during fermentation is through its known transcription factor interacting partners. At least two Med15 interacting TFs have a secondary role in silencing of subtelomeric genes. Oaf1 and Msn4 exert this effect by conditionally binding to telomeric proto-silencing regions known as X-elements without their usual co-activating protein partners, usually in a condition-specific manner. The negative regulatory function of Oaf1 corresponds to Oaf1 conditionally binding X elements at telomeres while growing in media with oleate (68). The WGJ environment may correspond to stabilization or increased binding of TFs to X-elements to promote silencing in a *MED15*-dependent manner. The reduced telomeric silencing in the deletion strain may be due to low levels of fatty acid production (a known phenotype of *med15* mutants) and thus less Oaf1 binding to X-elements.

### *MED15*_WY23_ Enhances Expression of Metabolic Genes Including Arginine Biosynthesis during Fermentation

We found previously that some *MED15* alleles from wine yeast confer enhanced fermentation traits when transplanted into lab strains (1). We therefore analyzed expression patterns in lab strains expressing one of two *MED15* alleles (WY7 and WY15) belonging to the European wine clade or a *MED15* allele from a palm wine yeast (WY23). A set of genes differentially regulated by WY alleles were identified by GO and pathway enrichment analyses (Fig. 6, 7, 8). *MED15*_WY23_ had the largest impact on the transcriptome. This allele differs from the LAB allele primarily in the length of poly-Q tracts, there being only 3 non-synonymous substitutions outside of poly-Q tracts, and no impact of removing them was observed in our previous studies (1). Expression of *MED15*_WY23_ in the lab strain background corresponded to a modest boost in alcoholic fermentation, suggesting that the observed gene expression differences may contribute to that phenotype. The differentially expressed genes in strains with *MED15*_WY23_ were enriched in metabolic activities, specifically amino acid biosynthesis including many arginine biosynthesis pathway enzymes (*ARG1*, 3-6) (Fig. 7). These same arginine biosynthesis pathway enzymes were all significantly differentially regulated in the *MED15*_LAB_ strain compared to a *med15* deletion strain (Fig. 4).

In the row normalized heat map shown in Fig. 9, it was possible to detect not only the previously noted significant decrease in *ARG1* and *ARG3*-*6* in the *med15* deletion mutant but also coordinated (albeit non-significant) decreases in *CPA1* and *CPA2* gene expression whose contribution to arginine biosynthesis can be seen in Fig. 4C, and *URA2*, a bifunctional protein whose N terminus comprises a carbamoyl phosphate synthetase domain that converts ammonium to carbamoyl-phosphate that feeds back into the urea cycle. Also apparent in the deletion mutant is a relative increase in expression of the *ARG2*, *ARG7* and *ARG8* genes that support the conversion of glutamate to ornithine, which feeds back into the arginine biosynthetic pathway via *ARG3*-dependent conversion to citrulline. The heat map also reveals that the *MED1*5_WY23_ allele is not the only WY allele affecting the relative expression of genes in the arginine biosynthetic pathway. Strains with *MED15*_WY7_ show a relative reduction in the same *ARG2*, *ARG7 ARG8* gene cluster that is more abundantly expressed in the deletion strain. These patterns suggest that the opposing expression of the two clusters of arginine biosynthetic genes is an important contributor to alcoholic fermentation, and that small *MED15* dependent adjustments that exaggerate the differential between the two clusters is one mechanism by which fermentation characteristics of a strain can be improved.

Arginine is one of the major amino acids found in grape must and an important nitrogen source for yeast. Arginine is cleaved by arginase (*CAR1* gene) into ornithine and urea, both of which can be used as a nitrogen source. Urea can be further catabolized to NH_3_ (ammonia) and CO_2_. This is an energy dependent process carried out by *DUR12* encoding urea amidolyase. Arginine and urea are considered secondary nitrogen sources and are used by the cell only when better ones like ammonia and glutamine are no longer present in the growth media. Constitutive expression of *DUR12* in wine yeast improves the degradation of urea during grape must fermentation, a reduction in fermentation time and increased ethanol production (69, 70). Arginine is also known to have a protective effect against ethanol stress in part by playing a role in the maintenance of cell wall integrity and in reducing oxidative stress (71) so elevated arginine levels in strains with *MED15*_WY_ alleles may also contribute to the ability of yeast strains to survive the stressful environment resulting from alcoholic fermentation.

### Arginine Biosynthesis Involvement in MED15-Dependent Fermentation Phenotypes

We tested the significance of arginine in the fermentation process by measuring the time to 50% weight loss in fermentations carried out in minimal (SD) media supplemented with different levels of arginine (Fig. 10B). Arginine supplementation increased fermentation efficiency regardless of the presence or of the specific *MED15* allele. In contrast, the impact of the *MED15* allele was clear in fermentations conducted in media lacking arginine. Strains with the *MED15*_WY7_ or *MED15*_WY23_ allele performed significantly better than strains with the *MED15*_LAB_ allele in arginine deficient media while strains with the *MED15*_WY15_ allele were unaffected (Fig. 10C). The *MED15*_WY7_ and *MED15*_WY23_ dependent transcriptomes facilitate arginine biosynthesis using two strategies; by promoting arginine production from glutamine (*CPA1*, *CPA2*, *ARG1,3, 4)* or by limiting ornithine production (Fig. 7, 9).

While *MED15*_WY7_ and *MED15*_WY15_ alleles had transcriptomes nearly identical to *MED15*_LAB_ at the tested time point, there were a small number of differentially expressed genes, and these overlapped with differentially expressed genes in the *MED15*_WY23_ analysis, including *MCH5*, *SNQ2*, and *LYS20*. Since these genes are differentially regulated by at least two of the tested *MED15*_WY_ alleles, they are also candidate contributors to alcoholic fermentation efficiency. *MCH5*, a riboflavin transport gene (72) was previously identified as a fermentation stress response gene induced early in fermentation (73). In addition to riboflavin in the environment, *MCH5* is regulated by proline (74), which may be relevant to the limited nitrogen environment of WGJ. *LYS20* encodes a homocitrate synthase and may be beneficial in the limited nitrogen environment of WGJ. Additionally, *LYS20* functions in DNA repair (75) which may also be beneficial in the harsh WGJ environment. *SNQ2* is a transporter involved in pleiotropic drug resistance regulated by Pdr1/3 (76), and has previously been shown to serve a protective role in the fermentation environment (77). In preliminary experiments, we found that deletion of the *SNQ2* gene in strains with the *MED15_WY7_* allele exhibited reduced fermentation (data not shown).

Analysis of pathways rather than GO enrichment terms led to additional hypotheses concerning gene expression patterns that may contribute to increased fermentation rates as summarized in Fig. 8. Roles for general amino acid biosynthesis, translation, DNA metabolism and mitochondrial dynamics in fine-tuning fermentation await further analysis.

One limitation of this study is that the influence of the *MED15*_WY_ alleles on gene expression may vary at different times during the fermentation process. Future transcriptomic analyses of these and additional strains throughout the time course of a fermentation reaction could reveal the full effect of *MED15* alleles on gene expression responsible for the boosted fermentation phenotypes we observed (1).

## Authors’ contributions

DC generated the data for Figures 1-9 and supplementary figures S1 and S2 and prepared the figures; EG generated the data for Figure 10. JF supervised the studies and contributed to the writing of the manuscript.

## Acknowledgements

We thank Dr. Cindy Toll for preparing libraries for RNA-Seq analysis, Dr. Daniel Weeks and Dr. Bryan Phillips for critical comments on this manuscript and the Iowa Institute of Human Genetics at the University of Iowa for generating the RNA-sequencing data.

## References

1. Cooper DG, Jiang Y, Skuodas S, Wang L, Fassler JS. Possible Role for Allelic Variation in Yeast MED15 in Ecological Adaptation. Front Microbiol. 2021;12:741572. doi:10.3389/fmicb.2021.741572.

2. Cooper DG, Fassler JS. Med15: Glutamine-Rich Mediator Subunit with Potential for Plasticity. Trends in biochemical sciences. 2019;44(9):737–51. doi:10.1016/j.tibs.2019.03.008.

3. Marsit S, Dequin S. Diversity and adaptive evolution of Saccharomyces wine yeast: a review. FEMS yeast research. 2015;15(7). doi:10.1093/femsyr/fov067.

4. Mortimer RK. Evolution and variation of the yeast (Saccharomyces) genome. Genome research. 2000;10(4):403–9. doi:10.1101/gr.10.4.403.

5. Fleet GH. Wine microbiology and biotechnology. Chur ; Philadelphia, Pa.: Harwood Academic Publishers; 1993. x, 510 p. p.

6. Pretorius IS. Tailoring wine yeast for the new millennium: novel approaches to the ancient art of winemaking. Yeast. 2000;16(8):675–729. doi:10.1002/1097-0061(20000615)16:8<675::AID-YEA585>3.0.CO;2-B.

7. Varela C, Torrea D, Schmidt SA, Ancin-Azpilicueta C, Henschke PA. Effect of oxygen and lipid supplementation on the volatile composition of chemically defined medium and Chardonnay wine fermented with Saccharomyces cerevisiae. Food chemistry. 2012;135(4):2863–71. doi:10.1016/j.foodchem.2012.06.127.

8. Peltier E, Bernard M, Trujillo M, Prodhomme D, Barbe JC, Gibon Y, et al. Wine yeast phenomics: A standardized fermentation method for assessing quantitative traits of Saccharomyces cerevisiae strains in enological conditions. PloS one. 2018;13(1):e0190094. doi:10.1371/journal.pone.0190094.

9. Gallone B, Steensels J, Prahl T, Soriaga L, Saels V, Herrera-Malaver B, et al. Domestication and Divergence of Saccharomyces cerevisiae Beer Yeasts. Cell. 2016;166(6):1397–410 e16. doi:10.1016/j.cell.2016.08.020.

10. Mendes-Ferreira A, del Olmo M, Garcia-Martinez J, Jimenez-Marti E, Leao C, Mendes-Faia A, et al. Saccharomyces cerevisiae signature genes for predicting nitrogen deficiency during alcoholic fermentation. Applied and environmental microbiology. 2007;73(16):5363–9. doi:10.1128/AEM.01029-07.

11. Mendes-Ferreira A, del Olmo M, Garcia-Martinez J, Jimenez-Marti E, Mendes-Faia A, Perez-Ortin JE, et al. Transcriptional response of Saccharomyces cerevisiae to different nitrogen concentrations during alcoholic fermentation. Applied and environmental microbiology. 2007;73(9):3049–60. doi:10.1128/AEM.02754-06.

12. Legras JL, Galeote V, Bigey F, Camarasa C, Marsit S, Nidelet T, et al. Adaptation of S. cerevisiae to Fermented Food Environments Reveals Remarkable Genome Plasticity and the Footprints of Domestication. Molecular biology and evolution. 2018;35(7):1712–27. doi:10.1093/molbev/msy066.

13. Johnston JR, Baccari C, Mortimer RK. Genotypic characterization of strains of commercial wine yeasts by tetrad analysis. Research in microbiology. 2000;151(7):583–90. doi:10.1016/s0923-2508(00)00228-x.

14. Fay JC, Benavides JA. Evidence for domesticated and wild populations of Saccharomyces cerevisiae. PLoS genetics. 2005;1(1):66–71. doi:10.1371/journal.pgen.0010005.

15. Crabtree HG. Observations on the carbohydrate metabolism of tumours. The Biochemical journal. 1929;23(3):536–45. doi:10.1042/bj0230536.

16. De Deken RH. The Crabtree effect: a regulatory system in yeast. J Gen Microbiol. 1966;44(2):149–56. doi:10.1099/00221287-44-2-149.

17. de Smidt O, du Preez JC, Albertyn J. The alcohol dehydrogenases of Saccharomyces cerevisiae: a comprehensive review. FEMS yeast research. 2008;8(7):967–78. doi:10.1111/j.1567-1364.2008.00387.x.

18. Wills C. Production of yeast alcohol dehydrogenase isoenzymes by selection. Nature. 1976;261(5555):26–9. doi:10.1038/261026a0.

19. Ihmels J, Bergmann S, Gerami-Nejad M, Yanai I, McClellan M, Berman J, et al. Rewiring of the yeast transcriptional network through the evolution of motif usage. Science. 2005;309(5736):938–40. doi:10.1126/science.1113833.

20. Gallagher JEG, Ser SL, Ayers MC, Nassif C, Pupo A. The Polymorphic PolyQ Tail Protein of the Mediator Complex, Med15, Regulates the Variable Response to Diverse Stresses. International journal of molecular sciences. 2020;21(5). doi:10.3390/ijms21051894.

21. Cooper DG, Liu S, Grunkemeyer E, Fassler JS. The Role of Med15 Sequence Features in Transcription Factor Interactions. Molecular and cellular biology. 2024:1–20. doi:10.1080/10985549.2024.2436672.

22. Gemayel R, Chavali S, Pougach K, Legendre M, Zhu B, Boeynaems S, et al. Variable Glutamine-Rich Repeats Modulate Transcription Factor Activity. Molecular cell. 2015;59(4):615–27. doi:10.1016/j.molcel.2015.07.003.

23. Bryan AC, Zhang J, Guo J, Ranjan P, Singan V, Barry K, et al. A Variable Polyglutamine Repeat Affects Subcellular Localization and Regulatory Activity of a Populus ANGUSTIFOLIA Protein. G3. 2018;8(8):2631–41. doi:10.1534/g3.118.200188.

24. O’Malley KG, Ford MJ, Hard JJ. Clock polymorphism in Pacific salmon: evidence for variable selection along a latitudinal gradient. Proceedings Biological sciences. 2010;277(1701):3703–14. doi:10.1098/rspb.2010.0762.

25. Caprioli M, Ambrosini R, Boncoraglio G, Gatti E, Romano A, Romano M, et al. Clock gene variation is associated with breeding phenology and maybe under directional selection in the migratory barn swallow. PloS one. 2012;7(4):e35140. doi:10.1371/journal.pone.0035140.

26. Mortimer RK, Johnston JR. Genealogy of principal strains of the yeast genetic stock center. Genetics. 1986;113(1):35–43.

27. Brachmann CB, Davies A, Cost GJ, Caputo E, Li J, Hieter P, et al. Designer deletion strains derived from Saccharomyces cerevisiae S288C: a useful set of strains and plasmids for PCR-mediated gene disruption and other applications. Yeast. 1998;14(2):115–32. doi:10.1002/(SICI)1097-0061(19980130)14:2<115::AID-YEA204>3.0.CO;2-2.

28. Bradbury JE, Richards KD, Niederer HA, Lee SA, Rod Dunbar P, Gardner RC. A homozygous diploid subset of commercial wine yeast strains. Antonie van Leeuwenhoek. 2006;89(1):27–37. doi:10.1007/s10482-005-9006-1.

29. Kim DH, Kim GS, Yun CH, Lee YC. Functional conservation of the glutamine-rich domains of yeast Gal11 and human SRC-1 in the transactivation of glucocorticoid receptor Tau 1 in Saccharomyces cerevisiae. Molecular and cellular biology. 2008;28(3):913–25. doi:10.1128/MCB.01140-07.

30. Ito H, Fukuda Y, Murata K, Kimura A. Transformation of intact yeast cells treated with alkali cations. Journal of bacteriology. 1983;153(1):163–8. doi:10.1128/JB.153.1.163-168.1983.

31. Gietz RD, Woods RA. Transformation of yeast by lithium acetate/single-stranded carrier DNA/polyethylene glycol method. Methods in enzymology. 2002;350:87–96. doi:10.1016/s0076-6879(02)50957-5.

32. Collart MA, Oliviero S. Preparation of yeast RNA. Current protocols in molecular biology. 2001;Chapter 13:Unit13 2. doi:10.1002/0471142727.mb1312s23.

33. Jalili V, Afgan E, Gu Q, Clements D, Blankenberg D, Goecks J, et al. The Galaxy platform for accessible, reproducible and collaborative biomedical analyses: 2020 update. Nucleic acids research. 2020;48(W1):W395–W402. doi:10.1093/nar/gkaa434.

34. Langmead B, Salzberg SL. Fast gapped-read alignment with Bowtie 2. Nature methods. 2012;9(4):357–9. doi:10.1038/nmeth.1923.

35. Liao Y, Smyth GK, Shi W. featureCounts: an efficient general purpose program for assigning sequence reads to genomic features. Bioinformatics. 2014;30(7):923–30. doi:10.1093/bioinformatics/btt656.

36. Love MI, Huber W, Anders S. Moderated estimation of fold change and dispersion for RNA-seq data with DESeq2. Genome biology. 2014;15(12):550. doi:10.1186/s13059-014-0550-8.

37. Goedhart J, Luijsterburg MS. VolcaNoseR is a web app for creating, exploring, labeling and sharing volcano plots. Scientific reports. 2020;10(1):20560. doi:10.1038/s41598-020-76603-3.

38. Hulsen T, de Vlieg J, Alkema W. BioVenn - a web application for the comparison and visualization of biological lists using area-proportional Venn diagrams. BMC genomics. 2008;9:488. doi:10.1186/1471-2164-9-488.

39. Balakrishnan R, Park J, Karra K, Hitz BC, Binkley G, Hong EL, et al. YeastMine--an integrated data warehouse for Saccharomyces cerevisiae data as a multipurpose tool-kit. Database : the journal of biological databases and curation. 2012;2012:bar062. doi:10.1093/database/bar062.

40. Chen EY, Tan CM, Kou Y, Duan Q, Wang Z, Meirelles GV, et al. Enrichr: interactive and collaborative HTML5 gene list enrichment analysis tool. BMC bioinformatics. 2013;14:128. doi:10.1186/1471-2105-14-128.

41. Kuleshov MV, Jones MR, Rouillard AD, Fernandez NF, Duan Q, Wang Z, et al. Enrichr: a comprehensive gene set enrichment analysis web server 2016 update. Nucleic acids research. 2016;44(W1):W90–7. doi:10.1093/nar/gkw377.

42. Xie Z, Bailey A, Kuleshov MV, Clarke DJB, Evangelista JE, Jenkins SL, et al. Gene Set Knowledge Discovery with Enrichr. Current protocols. 2021;1(3):e90. doi:10.1002/cpz1.90.

43. Karp PD, Billington R, Caspi R, Fulcher CA, Latendresse M, Kothari A, et al. The BioCyc collection of microbial genomes and metabolic pathways. Briefings in bioinformatics. 2019;20(4):1085–93. doi:10.1093/bib/bbx085.

44. Korotkevich G, Sukhov V, Budin N, Shpak B, Artyomov MN, Sergushichev A. Fast gene set enrichment analysis. 2021. doi:10.1101/060012.

45. Kamburov A, Herwig R. ConsensusPathDB 2022: molecular interactions update as a resource for network biology. Nucleic acids research. 2022;50(D1):D587–D95. doi:10.1093/nar/gkab1128.

46. Chen H, Boutros PC. VennDiagram: a package for the generation of highly-customizable Venn and Euler diagrams in R. BMC bioinformatics. 2011;12:35. doi:10.1186/1471-2105-12-35.

47. Kolde R. pheatmap: Pretty Heatmaps. 2025.

48. Hu Z, Killion PJ, Iyer VR. Genetic reconstruction of a functional transcriptional regulatory network. Nature genetics. 2007;39(5):683–7. doi:10.1038/ng2012.

49. Ansari SA, Ganapathi M, Benschop JJ, Holstege FC, Wade JT, Morse RH. Distinct role of Mediator tail module in regulation of SAGA-dependent, TATA-containing genes in yeast. The EMBO journal. 2012;31(1):44–57. doi:10.1038/emboj.2011.362.

50. Kemmeren P, Sameith K, van de Pasch LA, Benschop JJ, Lenstra TL, Margaritis T, et al. Large-scale genetic perturbations reveal regulatory networks and an abundance of gene-specific repressors. Cell. 2014;157(3):740–52. doi:10.1016/j.cell.2014.02.054.

51. Rossignol T, Dulau L, Julien A, Blondin B. Genome-wide monitoring of wine yeast gene expression during alcoholic fermentation. Yeast. 2003;20(16):1369–85. doi:10.1002/yea.1046.

52. Holt S, Miks MH, de Carvalho BT, Foulquie-Moreno MR, Thevelein JM. The molecular biology of fruity and floral aromas in beer and other alcoholic beverages. FEMS microbiology reviews. 2019;43(3):193–222. doi:10.1093/femsre/fuy041.

53. Pretorius I. Conducting Wine Symphonics with the Aid of Yeast Genomics. Beverages. 2016;2(4):36. doi:10.3390/beverages2040036.

54. Lenstra TL, Benschop JJ, Kim T, Schulze JM, Brabers NA, Margaritis T, et al. The specificity and topology of chromatin interaction pathways in yeast. Molecular cell. 2011;42(4):536–49. doi:10.1016/j.molcel.2011.03.026.

55. Hocher A, Ruault M, Kaferle P, Descrimes M, Garnier M, Morillon A, et al. Expanding heterochromatin reveals discrete subtelomeric domains delimited by chromatin landscape transitions. Genome research. 2018;28(12):1867–81. doi:10.1101/gr.236554.118.

56. Naumov G, Turakainen H, Naumova E, Aho S, Korhola M. A new family of polymorphic genes in Saccharomyces cerevisiae: alpha-galactosidase genes MEL1-MEL7. Molecular & general genetics : MGG. 1990;224(1):119–28. doi:10.1007/BF00259458.

57. Chow TH, Sollitti P, Marmur J. Structure of the multigene family of MAL loci in Saccharomyces. Molecular & general genetics : MGG. 1989;217(1):60–9. doi:10.1007/BF00330943.

58. Charron MJ, Read E, Haut SR, Michels CA. Molecular evolution of the telomere-associated MAL loci of Saccharomyces. Genetics. 1989;122(2):307–16.

59. Jiranek V, Langridge P, Henschke PA. Amino Acid and Ammonium Utilization by Saccharomyces cerevisiae Wine Yeasts From a Chemically Defined Medium. Am J Enol Vitic. 1995;46(1):75–83.

60. Xue T, Liu K, Chen D, Yuan X, Fang J, Yan H, et al. Improved bioethanol production using CRISPR/Cas9 to disrupt the ADH2 gene in Saccharomyces cerevisiae. World journal of microbiology & biotechnology. 2018;34(10):154. doi:10.1007/s11274-018-2518-4.

61. de Smidt O, du Preez JC, Albertyn J. Molecular and physiological aspects of alcohol dehydrogenases in the ethanol metabolism of Saccharomyces cerevisiae. FEMS yeast research. 2012;12(1):33–47. doi:10.1111/j.1567-1364.2011.00760.x.

62. Tamas MJ, Luyten K, Sutherland FC, Hernandez A, Albertyn J, Valadi H, et al. Fps1p controls the accumulation and release of the compatible solute glycerol in yeast osmoregulation. Molecular microbiology. 1999;31(4):1087–104. doi:10.1046/j.1365-2958.1999.01248.x.

63. Varelas V, Sotiropoulou E, Karambini X, Liouni M, Nerantzis E. Impact of Glucose Concentration and NaCl Osmotic Stress on Yeast Cell Wall β-d-Glucan Formation during Anaerobic Fermentation Process. Fermentation. 2017;3(3):44. doi:10.3390/fermentation3030044.

64. Kim KS, Yun HS. Production of soluble β-glucan from the cell wall of Saccharomyces cerevisiae. Enzyme and Microbial Technology. 2006;39(3):496–500. doi:10.1016/j.enzmictec.2005.12.020.

65. McMurrough I, Rose AH. Effect of growth rate and substrate limitation on the composition and structure of the cell wall of Saccharomyces cerevisiae. Biochemical Journal. 1967;105(1):189–203. doi:10.1042/bj1050189.

66. Ottaviani A, Gilson E, Magdinier F. Telomeric position effect: from the yeast paradigm to human pathologies? Biochimie. 2008;90(1):93–107. doi:10.1016/j.biochi.2007.07.022.

67. Peng J, Zhou JQ. The tail-module of yeast Mediator complex is required for telomere heterochromatin maintenance. Nucleic acids research. 2012;40(2):581–93. doi:10.1093/nar/gkr757.

68. Smith JJ, Miller LR, Kreisberg R, Vazquez L, Wan Y, Aitchison JD. Environment-responsive transcription factors bind subtelomeric elements and regulate gene silencing. Molecular systems biology. 2011;7:455. doi:10.1038/msb.2010.110.

69. Coulon J, Husnik JI, Inglis DL, van der Merwe GK, Lonvaud A, Erasmus DJ, et al. Metabolic Engineering ofSaccharomyces cerevisiaeto Minimize the Production of Ethyl Carbamate in Wine. American Journal of Enology and Viticulture. 2006;57(2):113–24. doi:10.5344/ajev.2006.57.2.113.

70. Shalamitskiy MY, Tanashchuk TN, Cherviak SN, Vasyagin EA, Ravin NV, Mardanov AV. Ethyl Carbamate in Fermented Food Products: Sources of Appearance, Hazards and Methods for Reducing Its Content. Foods. 2023;12(20). doi:10.3390/foods12203816.

71. Cheng Y, Du Z, Zhu H, Guo X, He X. Protective Effects of Arginine on Saccharomyces cerevisiae Against Ethanol Stress. Scientific reports. 2016;6:31311. doi:10.1038/srep31311.

72. Reihl P, Stolz J. The monocarboxylate transporter homolog Mch5p catalyzes riboflavin (vitamin B2) uptake in Saccharomyces cerevisiae. The Journal of biological chemistry. 2005;280(48):39809–17. doi:10.1074/jbc.M505002200.

73. Marks VD, Ho Sui SJ, Erasmus D, van der Merwe GK, Brumm J, Wasserman WW, et al. Dynamics of the yeast transcriptome during wine fermentation reveals a novel fermentation stress response. FEMS yeast research. 2008;8(1):35–52. doi:10.1111/j.1567-1364.2007.00338.x.

74. Spitzner A, Perzlmaier AF, Geillinger KE, Reihl P, Stolz J. The proline-dependent transcription factor Put3 regulates the expression of the riboflavin transporter MCH5 in Saccharomyces cerevisiae. Genetics. 2008;180(4):2007–17. doi:10.1534/genetics.108.094458.

75. Torres-Machorro AL, Aris JP, Pillus L. A moonlighting metabolic protein influences repair at DNA double-stranded breaks. Nucleic acids research. 2015;43(3):1646–58. doi:10.1093/nar/gku1405.

76. Rogers B, Decottignies A, Kolaczkowski M, Carvajal E, Balzi E, Goffeau A. The pleitropic drug ABC transporters from Saccharomyces cerevisiae. Journal of molecular microbiology and biotechnology. 2001;3(2):207–14.

77. Watanabe M, Mizoguchi H, Nishimura A. Disruption of the ABC transporter genes PDR5, YOR1, and SNQ2, and their participation in improved fermentative activity of a sake yeast mutant showing pleiotropic drug resistance. J Biosci Bioeng. 2000;89(6):569–76. doi:10.1016/s1389-1723(00)80059-6.

78. Sikorski RS, Hieter P. A system of shuttle vectors and yeast host strains designed for efficient manipulation of DNA in Saccharomyces cerevisiae. Genetics. 1989;122(1):19–27.

