## Supplementary Files for "Transcriptomic shift in ethanol and amino acid metabolic genes regulated by Med15 during alcoholic fermentation"

**^1^Department of Biology, University of Iowa, Iowa City, IA**

**^2^Department of Pharmaceutical Sciences, Butler University, Indianapolis, IN**

**^3^Corresponding Author**

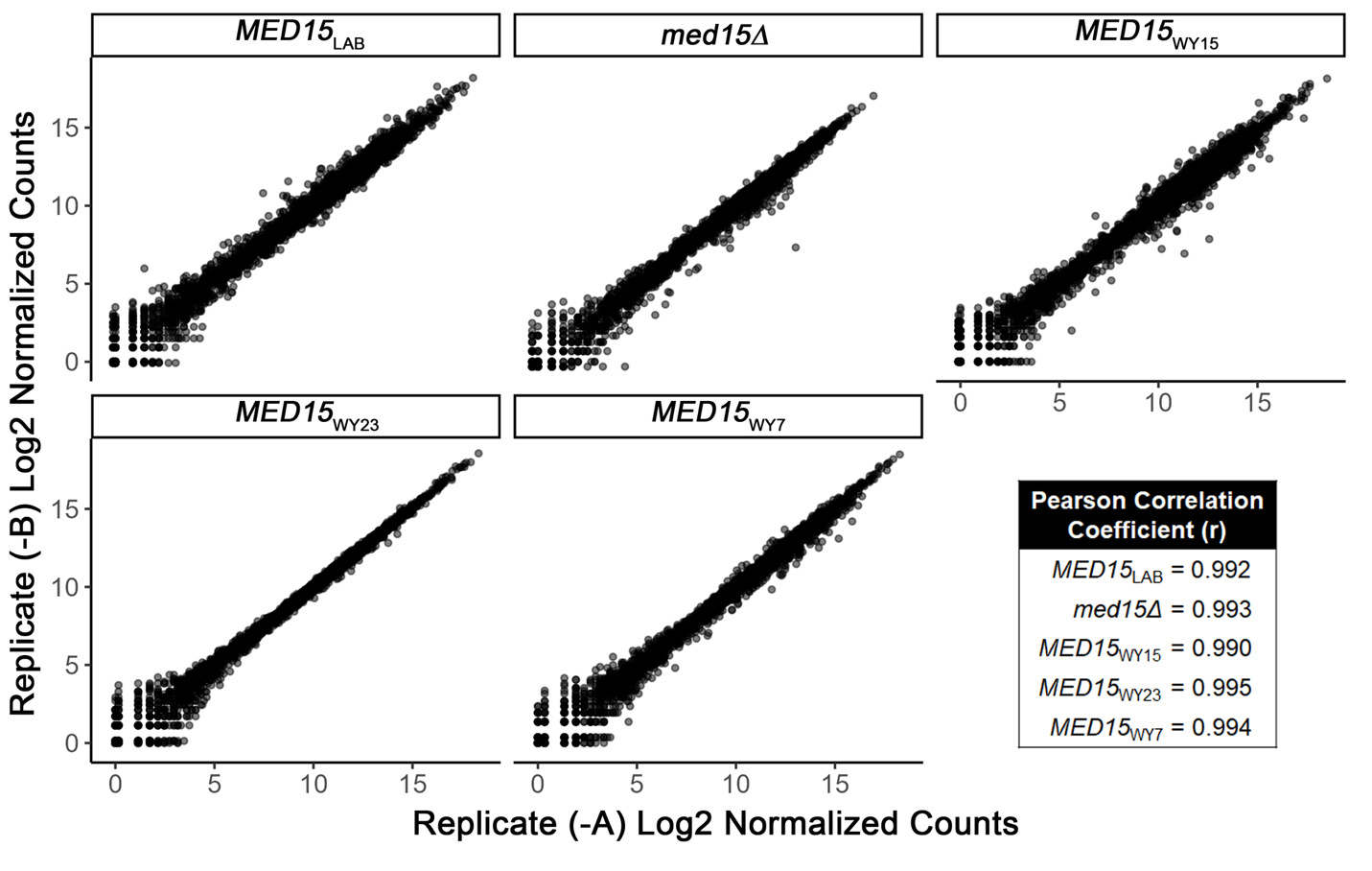

**Figure S1.** Correlation analysis between log2 transformed normalized counts of two biological replicates (-A replicate counts on x-axis and -B replicate counts on y-axis) for each *MED15* genotype. Pearson correlation coefficients (r) range between 0.990 and 0.995.

$`LAB:WY7:WY15`

```
[1] "Formation of a pool of free 40S subunits | Reactome"
[2] "GTP hydrolysis and joining of the 60S ribosomal subunit | Reactome"

$LAB
 [1] "Basal transcription factors - Saccharomyces cerevisiae (budding yeast) | KEGG"
 [2] "Estrogen-dependent gene expression | Reactome"
 [3] "Gene expression (Transcription) | Reactome"
 [4] "Glucose metabolism | Reactome"
 [5] "Glycolysis | Reactome"
 [6] "Metabolism of carbohydrates | Reactome"
 [7] "Orc1 removal from chromatin | Reactome"
 [8] "Pyruvate metabolism - Saccharomyces cerevisiae (budding yeast) | KEGG"
 [9] "RNA Polymerase II Pre-transcription Events | Reactome"
[10] "RNA Polymerase II Promoter Escape | Reactome"
[11] "RNA Polymerase II Transcription Elongation | Reactome"
[12] "RNA Polymerase II Transcription Initiation And Promoter Clearance | Reactome"
[13] "RNA Polymerase II Transcription Initiation | Reactome"
[14] "RNA Polymerase II Transcription Pre-Initiation And Promoter Opening | Reactome"
[15] "RNA Polymerase II Transcription | Reactome"
[16] "RNA polymerase II transcribes snRNA genes | Reactome"
[17] "Sulfur metabolism - Saccharomyces cerevisiae (budding yeast) | KEGG"
[18] "Switching of origins to a post-replicative state | Reactome"
[19] "gluconeogenesis | YeastCyc"

$WY7
 [1] "Base excision repair - Saccharomyces cerevisiae (budding yeast) | KEGG"
 [2] "Cysteine and methionine metabolism - Saccharomyces cerevisiae (budding yeast) | KEGG"
 [3] "DNA Double-Strand Break Repair | Reactome"
 [4] "Glycine, serine and threonine metabolism - Saccharomyces cerevisiae (budding yeast) | KEGG"
 [5] "Homologous recombination - Saccharomyces cerevisiae (budding yeast) | KEGG"
 [6] "MAPK signaling pathway - yeast - Saccharomyces cerevisiae (budding yeast) | KEGG"
 [7] "Metabolism | Reactome"
 [8] "Mismatch Repair | Reactome"
 [9] "Mismatch repair (MMR) directed by MSH2:MSH6 (MutSalpha) | Reactome"
[10] "Phenylalanine, tyrosine and tryptophan biosynthesis - Saccharomyces cerevisiae (budding yeast) | KEGG"
[11] "RNA Polymerase I Promoter Clearance | Reactome"
[12] "RNA Polymerase I Promoter Escape | Reactome"
[13] "RNA Polymerase I Transcription | Reactome"
[14] "SUMOylation of DNA damage response and repair proteins | Reactome"
[15] "cysteine biosynthesis IV (fungi) | YeastCyc"
[16] "superpathway of phenylalanine, tyrosine and tryptophan biosynthesis | YeastCyc"
[17] "superpathway of sulfur amino acid biosynthesis (<i>Saccharomyces cerevisiae</i>) | YeastCyc"

$WY15
[1] "Metabolism of proteins | Reactome"
[2] "TCA cycle II (plants and fungi) | YeastCyc"

$WY23
[1] "L-lysine biosynthesis IV | YeastCyc"
[2] "Respiratory electron transport, ATP synthesis by chemiosmotic coupling, and heat production by uncoupling proteins. | Reactome"
[3] "aerobic respiration (cytochrome c) | YeastCyc"
[4] "aerobic respiration (linear view) | YeastCyc"
[5] "urea cycle | YeastCyc"

$`LAB:WY7`
 [1] "Activation of the pre-replicative complex | Reactome"
 [2] "Cell Cycle | Reactome"
 [3] "Cell Cycle, Mitotic | Reactome"
 [4] "Cell cycle - yeast - Saccharomyces cerevisiae (budding yeast) | KEGG"
 [5] "DNA Replication Pre-Initiation | Reactome"
 [6] "DNA Replication | Reactome"
 [7] "DNA replication - Saccharomyces cerevisiae (budding yeast) | KEGG"
 [8] "DNA strand elongation | Reactome"
 [9] "Establishment of Sister Chromatid Cohesion | Reactome"
[10] "G1/S Transition | Reactome"
[11] "Glycolysis / Gluconeogenesis - Saccharomyces cerevisiae (budding yeast) | KEGG"
[12] "Lagging Strand Synthesis | Reactome"
[13] "Meiosis - yeast - Saccharomyces cerevisiae (budding yeast) | KEGG"
[14] "Mismatch repair - Saccharomyces cerevisiae (budding yeast) | KEGG"
[15] "Mitotic G1 phase and G1/S transition | Reactome"
[16] "Mitotic Prometaphase | Reactome"
[17] "Oxidative phosphorylation - Saccharomyces cerevisiae (budding yeast) | KEGG"
[18] "Processive synthesis on the lagging strand | Reactome"
[19] "Removal of the Flap Intermediate | Reactome"
[20] "Resolution of Sister Chromatid Cohesion | Reactome"
[21] "S Phase | Reactome"
[22] "Synthesis of DNA | Reactome"
[23] "Valine, leucine and isoleucine biosynthesis - Saccharomyces cerevisiae (budding yeast) | KEGG"
[24] "glycolysis III (from glucose) | YeastCyc"
[25] "glycolysis | YeastCyc"

$`LAB:WY23`
[1] "arginine biosynthesis | YeastCyc"

$`WY7:WY15`
 [1] "Activation of the mRNA upon binding of the cap-binding complex and eIFs, and subsequent binding to 43S | Reactome"
 [2] "Cap-dependent Translation Initiation | Reactome"
 [3] "Eukaryotic Translation Initiation | Reactome"
 [4] "Formation of the ternary complex, and subsequently, the 43S complex | Reactome"
 [5] "L13a-mediated translational silencing of Ceruloplasmin expression | Reactome"
 [6] "Metabolism of RNA | Reactome"
 [7] "Nonsense Mediated Decay (NMD) enhanced by the Exon Junction Complex (EJC) | Reactome"
 [8] "Nonsense Mediated Decay (NMD) independent of the Exon Junction Complex (EJC) | Reactome"
 [9] "Nonsense-Mediated Decay (NMD) | Reactome"
[10] "Ribosomal scanning and start codon recognition | Reactome"
[11] "Ribosome - Saccharomyces cerevisiae (budding yeast) | KEGG"
[12] "SRP-dependent cotranslational protein targeting to membrane | Reactome"
[13] "Translation initiation complex formation | Reactome"
[14] "Translation | Reactome"

$`WY7:WY23`
[1] "Lysine biosynthesis - Saccharomyces cerevisiae (budding yeast) | KEGG"
[2] "superpathway of leucine, valine, and isoleucine biosynthesis | YeastCyc"

$`WY15:WY23`
[1] "Citrate cycle (TCA cycle) - Saccharomyces cerevisiae (budding yeast) | KEGG"
[2] "Citric acid cycle (TCA cycle) | Reactome"
[3] "Pyruvate metabolism and Citric Acid (TCA) cycle | Reactome"
[4] "The citric acid (TCA) cycle and respiratory electron transport | Reactome"
```

**Figure S2.** Raw Output for Pathway Analysis corresponding to Venn Diagram (Fig. 8).

**Table S1. Genes differentially regulated by *MED15*_WY23_ during WGJ fermentation**

| **Higher expression in *MED15*_WY23_ strain** | | |
| --- | --- | --- |
| YER062C | GPP2 | Glycerol-3-Phosphate Phosphatase |
| YER069W | ARG5,6 | ARGinine requiring |
| YER073W | ALD5 | ALdehyde Dehydrogenase |
| YHR018C | ARG4 | ARGinine requiring |
| YKL029C | MAE1 | MAlic Enzyme |
| YKL120W | OAC1 | OxaloAcetate Carrier |
| YDL182W | LYS20 | LYSine requiring |
| YDR011W | SNQ2 | Sensitivity to 4-NitroQuinoline-N-oxide |
| YDR035W | ARO3 | AROmatic amino acid requiring |
| YDR158W | HOM2 | HOMoserine requiring |
| YGL009C | LEU1 | LEUcine biosynthesis |
| YGL184C | STR3 | Sulfur TRansfer |
| YJL088W | ARG3 | ARGinine requiring |
| YJR111C | PXP2 | PeroXisomal Protein |
| YML130C | ERO1 | ER Oxidation or Endoplasmic Reticulum Oxidoreductin |
| YOL058W | ARG1 | ARGinine requiring |
| YOR107W | RGS2 | Regulator of heterotrimeric G protein Signaling |
| YOR204W | DED1 | Defines Essential Domain |
| YOR267C | HRK1 | Hygromycin Resistance Kinase |
| YOR302W |  |  |
| YOR303W | CPA1 | Carbamyl Phosphate synthetase A |
| YOR306C | MCH5 | MonoCarboxylate transporter Homolog |
| YOR337W | TEA1 | Ty Enhancer Activator |
| YPL092W | SSU1 | Sensitive to SUlfite |
| **Lower expression in WY23 *MED15* strain** | | |
| YKL096W | CWP1 | Cell Wall Protein |
| YKL109W | HAP4 | Heme Activator Protein |
| YDR342C | HXT7 | HeXose Transporter |
| YDR406W | PDR15 | Pleiotropic Drug Resistance |
| YNL194C |  |  |
| YOR028C | CIN5 | Chromosome INstability |
| YPL230W | USV1 | Up in StarVation |
